# Does sequence clustering confound AlphaFold2?

**DOI:** 10.1101/2024.07.29.605333

**Authors:** Hannah K. Wayment-Steele, Sergey Ovchinnikov, Lucy Colwell, Dorothee Kern

## Abstract

Predicting multiple conformational states of proteins represents a significant open challenge in structural biology. Increasingly many methods have been reported for perturbing and sampling AlphaFold2 (AF2) [1] to achieve multiple conformational states. However, if multiple methods achieve similar results, that does not in itself invalidate any method, nor does it answer *why* these methods work. Interpreting why deep learning models give the results they do is a critically important endeavor for future model development and appropriate usage. To help the field continue to try to answer these questions, this work addresses misunderstandings and inaccurate conclusions in refs. [2-6]. Deep learning methods development moves quickly, and by no means did we think that the implementation of AF-Cluster in [7] would be the final word on how to sample multiple conformations. However, Porter et al.’s primary critique, that AF-Cluster does not use local evolutionary couplings in its MSA clusters, is incorrect. We report here further analysis that underscores our original finding that local evolutionary couplings do indeed play an important role in AF-Cluster predictions, and refute all false claims made against [7].

## Introduction

Proteins interconvert between multiple conformational states that are critical for their biological function. Predicting these alternative conformations from sequence alone has been a persistent challenge in computational structural biology. The advent of AlphaFold2 (AF2) has revolutionized protein structure prediction, but its application to proteins with multiple conformational states remains an active area of research. Because AF2 was trained with both sequence and structure information, we expect that sequence and structure information are convolved, and disentangling the effect of either is difficult. AF2 was trained with the task to predict the most likely structure given its training data, and if a protein appears in state A ten times as often as it appears in state B in the training data, it will favor predicting state A, regardless of the Boltzmann weights that state A and B might actually have in solution. For instance, ref. [8] compared ligand-binding proteins that had more apo or holo structures in the PDB, and found that random MSA subsampling in AF2 returned the apo state more for proteins with more solved apo structures, and the holo state more for proteins with more solved holo structures.

Our work introduced AF-Cluster [7], a method that leverages sequence clustering of multiple sequence alignments (MSAs) to predict multiple conformational states within protein families. The method revealed that different regions of sequence space within a protein family can exhibit preferences for different structural states, as demonstrated in fold-switching proteins like KaiB, RfaH, and Mad2. In developing AF-Cluster, we had focused on what could be achieved by modulating the sequence input for a system for which other methods focusing on randomly restricting the number of sequences input into AF2 MSA were unable to predict multiple conformations [9-11].

Why might the family of a metamorphic protein contain pockets of evolution surrounding both known folded states? KaiB and RfaH are two well-studied metamorphic proteins with very different biological roles – KaiB is involved in regulating bacterial circadian rhythms, whereas RfaH is a transcription factor – but both are posited to have evolved via gene duplication from proteins that did not switch folds. Both therefore are posited to have developed metamorphic behavior as a regulatory strategy.

Cyanobacteria contain up to 4 gene copies of KaiB [12]. KaiB-1, which is the best studied and the metamorphic protein that regulates circadian rhythms, has been proposed to evolve from a thioredoxin-like ancestor, acquiring the ability to switch into a binding-incompetent fold as a means of regulating its interaction with KaiC to regulate circadian rhythms [13]. Supporting this is the existence of modern KaiB variants that are stabilized in a thioredoxin-like conformation [7], such as that from L. pneumophilia [14], which is not involved in circadian rhythm activity.

RfaH arose via gene duplication from the general transcription elongation factor NusG, whose C-terminal β-barrel KOW domain binds the ribosomal protein S10 and promotes Rho-dependent termination [15, 16]. RfaH retained the ancestral NGN domain involved in RNA polymerase interactions, but evolved an autoinhibited state in which the KOW domain refolds to an α-helical hairpin [15]. This state sterically occludes the RNA polymerase interaction interface, and precludes premature binding to either S10 or RNA polymerase. Upon encountering an ops DNA element, RfaH refolds into a NusG-like β-barrel, restoring its binding capabilities.

Therefore, fold-switching appears to have emerged as a regulatory mechanism via gene duplication in both KaiB and RfaH. Supporting this concept in a different system, Volkman and colleagues demonstrated how the metamorphic protein Lymphotactin could evolve from a family with a single fold using ancestral sequence reconstruction [17] The premise of sequence clustering reflects what is increasingly understood about metamorphic families: that evolution could drive separation of distinct sequence groups within these families that evolve around one or the other conformational state. Indeed, the computational method by Porter and colleagues in ref. [18] also uses sub-families of MSAs as input to models for predicting multiple sets of contacts.

AF-Cluster has been subject to critique in 5 recent publications solely from the Porter group (3 preprints and 2 peer-reviewed) [2-6]. We present our responses to critiques of AF-Cluster in chronological order, finishing with the most pressing question to arise from these critiques in the preprint / Matters Arising titled “Sequence Clustering Confounds AlphaFold2” [5,6], which is whether evolutionary coupling information plays a role in AF-Cluster’s ability to predict alternative conformations and whether simpler approaches might achieve similar results. To respond to this point, we first contextualize how two different types of analyses we performed in our paper – analyzing sequence preferences over a protein family or over individual family members – got confused and led to misunderstandings in [5,6]. We next describe why the constructed CF-Random method in refs. [5,6] is not random sampling: it combines multiple AF2 settings that still must be individually (subjectively) selected and is not a direct test for evolutionary couplings in AF-Cluster. We conclude with a new direct test for the role of evolutionary couplings by comparing predictions from MSA clusters to those from MSAs with shuffled columns, which ablates coevolutionary signals while preserving single-site conservation. Our results demonstrate that evolutionary information embedded in MSA clusters is indeed used by AF2 to predict multiple conformational states, thereby clarifying this important question of the role of coevolutionary signal in modern structure prediction for the field.

### Response to “Colabfold predicts alternative protein structures from single sequences, coevolution unnecessary for AF-cluster”[2]

In the preprint in [2], the authors note that for 7 KaiB variants, using a single sequence enables ColabFold [19] to predict a structure known experimentally to be thermodynamically most stable. This confirms our analysis that was presented in Supplemental Discussion Figure 1d in our original paper [7] (reproduced here in **Figure 1A**), which shows that for 71% of the 487 variants examined, the single-sequence predictions match the predictions made using “shallow” MSAs. We are pleased that Porter et al. were able to reproduce a subset of these results. *Crucially, however, for roughly 50% of structures predicted to be in the FS state, predictions from single sequences differed compared to shallow MSAs (Figure 1a from our original paper* [7], *an observation that is not explained by Porter et al.’s analysis in* [2].

**Figure 1.**
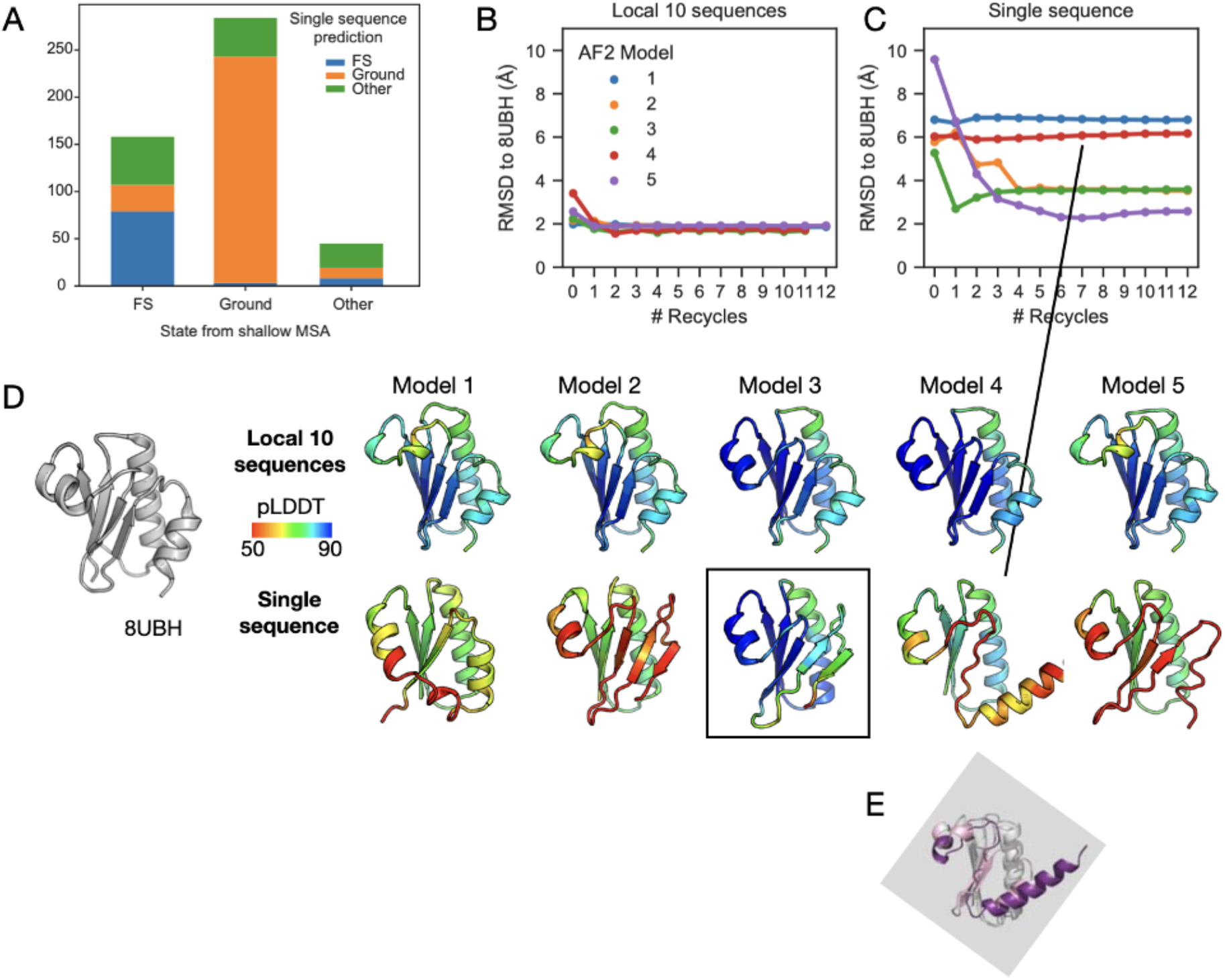
A) Reproduced supplemental discussion Figure 1d from [7]. Comparing models of KaiB variants predicted using shallow MSAs and single sequences. For variants predicted in the FS state using shallow MSAs, roughly only 50% are predicted in the FS state using single sequences. (B) Examining all 5 models and the effect of multiple recycles, a 10-sequence MSA creates significantly more robust, and correct predictions for KaiB^TV^-4 than a single sequence (C). D) Comparing our NMR structure of KaiB^TV^-4 (8UBH) [7] with structural models generated with the “local 10” sequence MSA (top) or single-sequence mode (bottom) at 7 recycles. Models were selected at the recycle at which any model from single-sequence mode obtains the lowest RMSD to our solved NMR structure. Model 3 (boxed) with highest pLDDT for single sequence mode as argued by Porter et al. in [2] predicts the wrong structure. E) Picked structure of KaiB^TV^-4 from Fig. 1 in [2] from single sequence mode, which was used by Porter et al. to claim that single sequence predicts correct structures, rotated to the same orientation, is the incorrect structure, compared to our solved NMR structure of KaiB^TV^-4 [7], invalidating claims made in Fig. 1 in [2].

[2] refers to our KaiB variant predictions using the 10 closest sequences from our phylogenetic tree by edit distance as “AF-Cluster” predictions, which is not how we defined the “AF-Cluster” method: In [7], we wrote, “From here on we refer to this entire pipeline as “AF-Cluster” – generating a MSA with ColabFold, clustering MSA sequences with DBSCAN, and running AF2 predictions for each cluster.” This method results *in MSAs of hugely different sizes*. We want to emphasize here that how subsampled MSAs should optimally be constructed is an interesting question.

We provide additional analysis buttressing our original conclusion that coevolutionary signal did play a role in AF2 structure prediction from a MSA of the closest 10 sequences for the construct KaiB^TV^-4, which formed a direct experimental test for our predictions and is one of the seven KaiB variants for which Porter et al. claimed co-evolutionary information was not needed [2]. To be consistent with our original wording in [7], we refer to MSAs constructed of the 10 closest sequences from the phylogenetic tree as “shallow” MSAs. We compared predictions from the shallow MSA and no MSA. We find that with the shallow MSA, all models converge within 1 recycle to 2 Å of the NMR structure with high confidence (**Figure 1B,D** upper row). In contrast, with a single sequence, *4 models result in wrong structures* even after many recycles, and only model 5 obtains the lowest RMSD to 8UBH (2.28 Å) in 7 recycles (**Figure 1C,D** lower row). The structure from Model 5 has attained approximately the correct fold (last helix and b-strand not quite formed), but with low confidence throughout the fold-switching region. Without prior knowledge, from the output of the single sequence predictions, one might pick the incorrect structural prediction, model 3, as it has the highest confidence. In contrast, the structures predicted using the local MSA have all converged to the correct fold and with high confidence, highlighting the improved performance of shallow MSAs over no MSA.

Furthermore, we recently showed [13] that single-sequence mode in ColabFold does not predict the actual Ground state for KaiB from *Rhodobacter sphaeroides*, one of the seven investigated in [2], but rather a register-shifted alternate conformation populated to about 6% at 20°C which we termed the “Enigma” state [13].

### Response to “AlphaFold2 has more to learn about protein energy landscapes” [3], later published in [4]

The premise of AF-Cluster was that a single protein family can contain clustered differing sequence preferences for more than one structure, and that by clustering the MSA input, AF2 is capable of detecting these preferences. [3] claims that AF-Cluster does not predict multiple conformations for a pair of two protein isoforms, BCCIP-alpha and BCCIP-beta [20]. However, this is not an example where we would expect AF-Cluster to be applicable. These two isoforms have completely different sequences for the last ∼20% of the protein due to alternative splicing. Constructing an MSA for BCCIP-alpha shows that ColabFold does not identify any sequence coverage for the region where the sequences differ (**Appendix A**). Therefore, no coevolutionary information exists from the outset in the MSA for AF2 or AF-Cluster to use to distinguish differing sequence preferences.

The second example given in [3] is an engineered fold-switching protein, S_A1[21]_. Since it is engineered, we do not expect the principle underlying AF-Cluster to apply – namely, the principle that natural protein families have evolved to contain more than one structure preference.

### Response to “Sequence clustering confounds AlphaFold2” on bioRxiv [5]

The commentary in the preprint “Sequence clustering confounds AlphaFold2” [5] can be grouped into two themes. The first, that AF-Cluster is a “poor predictor of metamorphic proteins”, suggests a fundamental misunderstanding of the method, which we discuss below. The second theme is that our original paper contained calculations that Schafer et al. failed to reproduce. However, the calculations on RfaH presented in [5] use different, older AF2 settings and therefore cannot be directly compared with results in our paper (see **Appendix B**). These false claims of “missing controls” were consequently taken out during the review process with Nature for the Matter Arising [6] in response to our formal written response and the reviewers evaluation.

The critique that AF-Cluster is a “poor predictor of metamorphic proteins” [5] misunderstands the method – AF-Cluster was developed as a method to detect the distribution of structure preferences across an entire protein family. This is encapsulated in **Figure 2A**, reproduced from [7]. A key finding we conveyed was that clusters of sequences predicted both low-and high-confidence structures across different regions of sequence space. We found this to be an intriguing “feature” rather than a “bug”, and further investigated this in our paper by creating a phylogenetic tree and exploring sequence-specific predictions using a different computational protocol than AF-Cluster (**Figure 2B**, reproduced from [7]). The observation in Fig. 1c in ref. [5] and [6] that AF-Cluster predicts multiple states for fold-switching proteins known to not occupy more than one state is missing the heart of the model.

**Figure 2.**
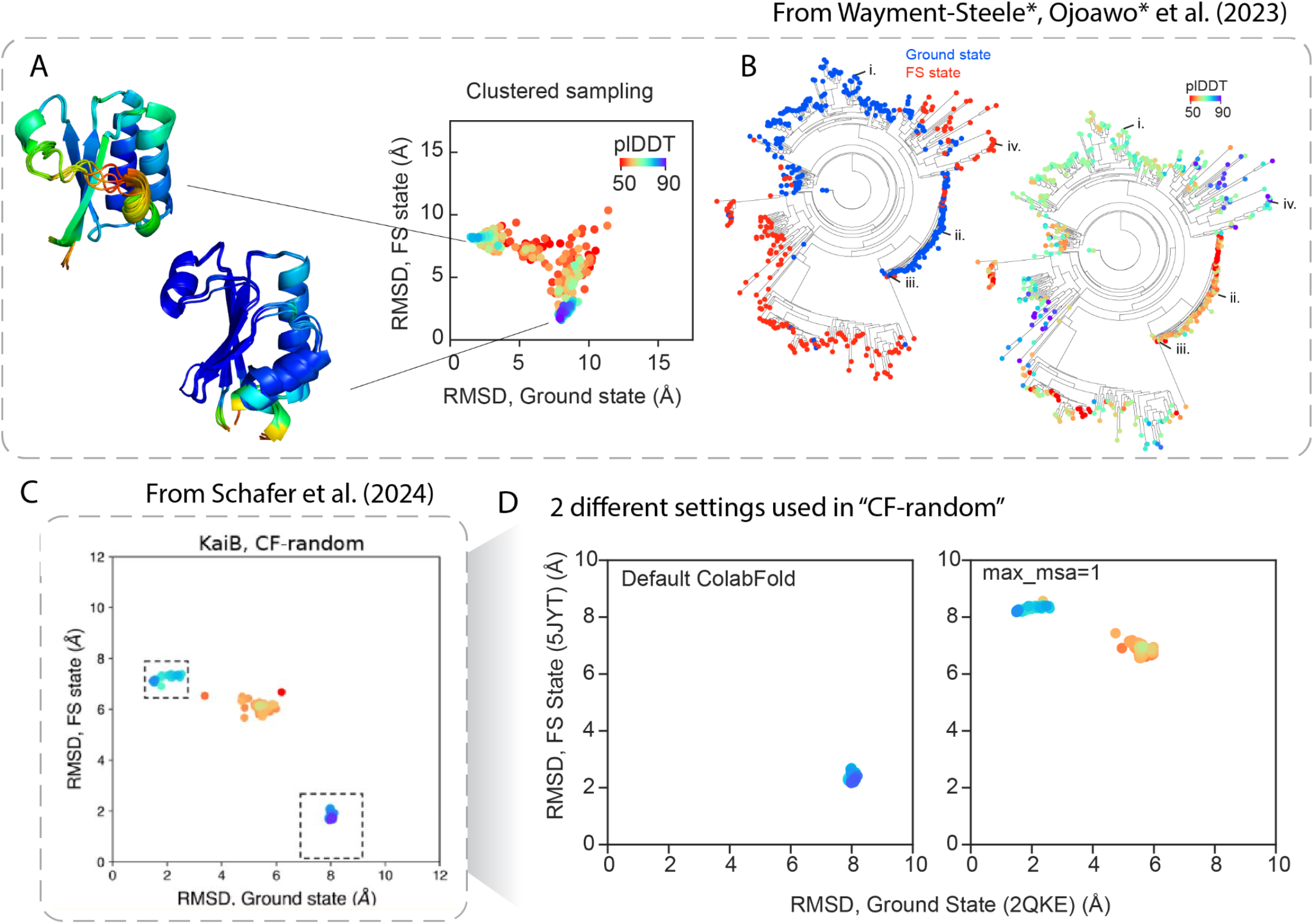
A) Reproduced from [7], Figure 1e/f, demonstrating that clustering the KaiB multiple sequence alignment (MSA) results in a distribution of structures, where the highest confidence structures are the two known well-populated states of KaiB. B) Reproduced from Figure 2a in [7], highlighting our investigation into KaiB sequence preferences to reveal that AF2 predicts strong sequence preferences across the family for both states in clusters across the family. C) Reproduced from Figure 2a in [5], claiming CF-random more efficiently predicts both states with high confidence. D) Separating the two different settings used in CF-random in [5] demonstrates that the two settings selected uniquely predict only one or the other state, as we had already reported in [7].

[5] argues that high-confidence parts of the landscape of structure preferences can be recapitulated using “CF-Random” (**Figure 2C**, reproduced from [5]). This method was presented as a route of using random sampling in ColabFold to predict multiple states. The authors description of this method [6], “we ran ColabFold -an efficient-yet-accurate implementation of AF2-with randomly sampled shallow input MSAs … This approach, hereafter called CF-random” obscures the fact that this method cherry-picks two different settings in ColabFold that each predict one of KaiB’s two states (**Figure 2D**); importantly, these settings simply reproduce results reported in our original paper. We already established in [7] that the full MSA for KaiB predicts the Fold-switched (FS) state, and we observe that in CF-random, all the FS state structures come from the run using the full MSA, while all ground state structures use the setting max-msa=1:2. The “1” in this setting means that one single sequence is selected to use as the MSA. AF2 always includes the original query sequence when it is performing this random sampling, so when just one sequence is selected, this results in always using the query sequence as the MSA. The “2” means that two sequences are randomly selected to use in the ‘extra_msa’ track. We also established that many variants, in single sequence mode, predict the Ground state, though not all (see response above to [2]).

CF-Random in preprint form [5] used different settings for each protein family presented, underscoring that the “CF-Random” method presented in [5] does not generalize. When this preprint was published as a Matters Arising [6], this part of the framework was modified as a result to our peer-reviewed response to the original Matter Arising version. Critically, CF-Random as published in [6] includes comparing to known structures in the PDB to pick an alternate conformation, which again would not generalize for proteins with uncharacterized alternate states. In contrast, AF-Cluster is an automated method using the statistical method DBSCAN, and is not dependent on current information on known structures.

Notably, Fig. 1b in [5], which compares CF-Random runs to sequence-specific predictions in [7], makes misleading comparisons in terms of sampling (**Figure 3**). The models from CF-Random that are shown in Fig. 1b are selected as max. pLDDT models from all sampled structures. In contrast, the RMSD and pLDDT comparisons that Schafer et al. used to compare for AF-cluster models were models from the AF-Cluster data repository, which used just 1 AF2 model and 1 random seed for each MSA cluster. Schafer et al. did not compare the two methods with equivalent sampling. As a representative example, if we sample from the MSA cluster for KaiB^RS^ using the same amount of sampling used in CF-Random (5 AF2 models, 5 seeds) [6], the maximum pLDDT sampled increases from 63.8 to 70.8 (Fig. 3). We also could not exactly reproduce the top model reported in CF-Random data repository with CF-Random’s reported settings in [6].

**Figure 3.**
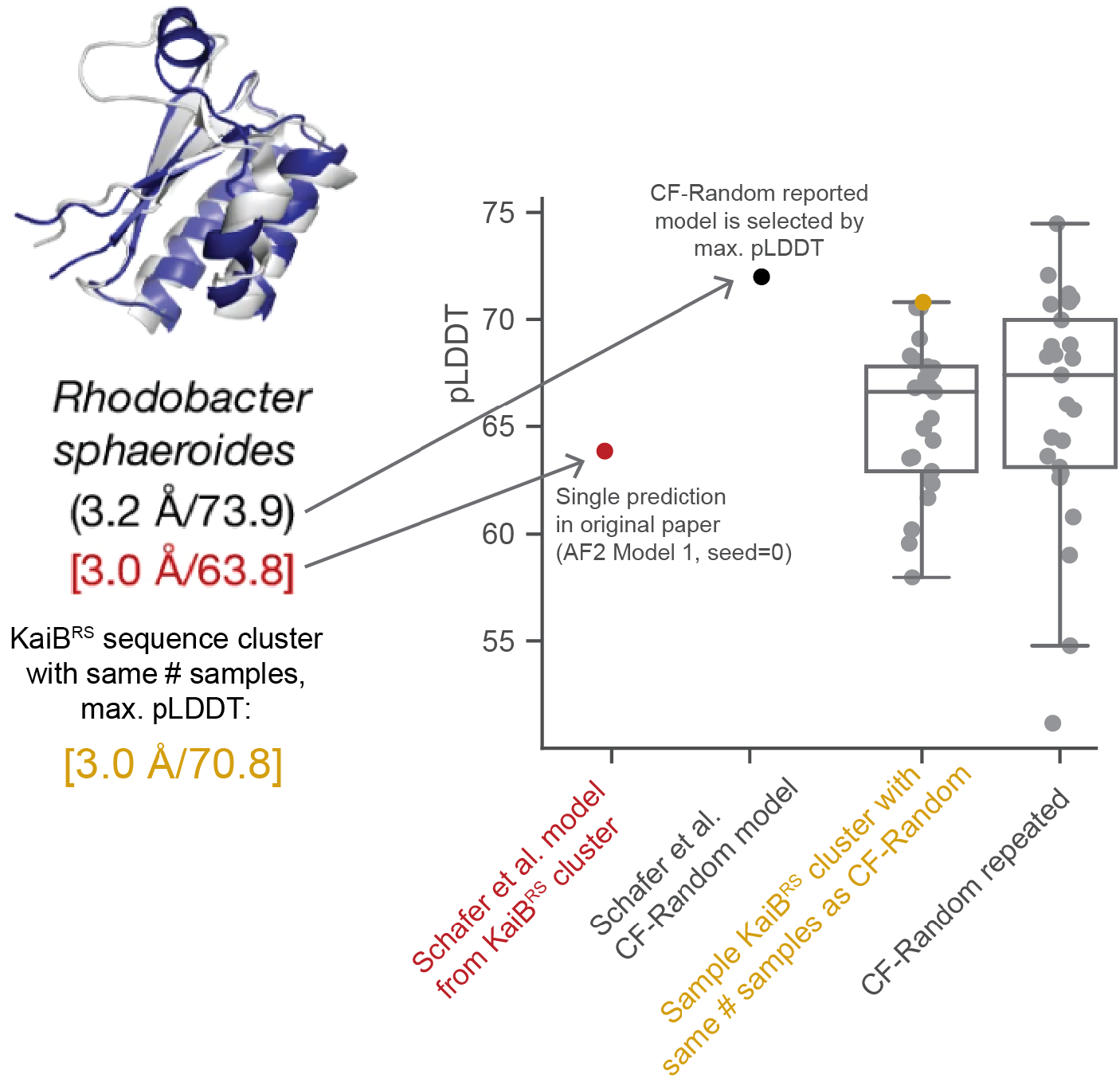
Ref. [6] Figure 1b makes misleading comparisons by selecting models from different amounts of sampling. CF-Random models are selected by max. pLDDT, and the AF-Cluster models used in their analysis were not. Repeating AF-Cluster sampling for the sequence cluster corresponding to KaiBRS with the same number of seeds that was used for CF-Random results in a highest-pLDDT model that goes from 63.8 to 70.8. We emphasize we do not view maximizing pLDDT as the primary objective of AF-Cluster – we found it quite interesting to find a distribution of pLDDTs sampled across the KaiB family (cf. Figure 2B).

Again, we emphasize that we do not view obtaining high pLDDT as a main parameter to optimize --we originally were intrigued by the range of pLDDTs reported across the KaiB family. We also reiterate what was originally stated in our paper [7]: “Firstly, the pLDDT metric itself cannot be used as a measure of free energy. This was immediately evident in our investigation of KaiB, where in our models generated with AF-Cluster, the thermodynamically-disfavored FS state, still had higher pLDDT than the ground state”. Finding better scoring metrics that more accurately reflect free energies or other desired metrics for optimization will aid the field significantly. Metrics based on other outputs from AF2 such as ipTM or PAE and which more explicitly account for low pLDDT in disordered regions may already provide better insight [22].

We also find insufficient evidence for [5]’s claim that MSA Transformer predicts contacts unique to the opposite structure from what AF-Cluster predicts (Appendix C). Such an analysis ought to be carried out across many clusters, and not just one, similarly to how we analyzed many clusters for KaiB in [7] that demonstrated coevolutionary contact predictions for each state (Extended Data Fig. 3 in [7]). We analyzed many clusters from RfaH as input into MSA Transformer and compared predicted contacts from the clusters that predicted either state in [7]. When comparing contact maps from MSA Transformer across multiple MSA clusters, we find that contacts unique to each state can indeed be identified (see Appendix C).

Finally, [5]’s claim that CF-Random is more efficient is also incorrect: when wall time is correctly tallied, CF-Random as reported and AF-Cluster as reported use equivalent sampling. We again emphasize, however, that we do not expect that every cluster should return a high-pLDDT structure, making high-pLDDT return rate an inappropriate measure of success for AF-Cluster. Indeed, the authors’ claim in [2] does not take pLDDT into account.

### Response to “Sequence clustering confounds AlphaFold2” published as Matters Arising [6]

The Matters Arising publication is titled the same as [5] on bioRxiv, and at the start of peer review contained the same arguments as [5]. During peer review, a new analysis was added by the Porter lab to try to support the claim that “Evolutionary couplings do not drive AF2-based predictions of alternative protein conformations”, including AF-Cluster. We submitted a formal response that was sent to reviewers, who recommended publication of the original Matters Arising together with our original response. However, we were not given a chance to see the final version of the Matters Arising or submit of a final response. Below is our response to the new argument by Porter eta al. that was added during peer review.

Schafer et al. argue that because no evolutionary couplings can be detected in the shallow MSAs used as part of CF-Random, it cannot be the case that AF2 [1] is using evolutionary coupling information from sequence clusters in AF-Cluster. This logic does not hold. Firstly, this conclusion disregards a study performed in [7] that demonstrated that the same MSA clusters, if input into MSA Transformer, predict contacts that trend with the predicted structures from AF2 across all the clusters (Extended Data Fig. 3 in [7]). As it is increasingly clear that there are many factors influencing AlphaFold predictions, to directly make claims about the influence of any given factor such as evolutionary couplings, the field needs interpretability tests that directly query specific factors of interest. Constructing the “CF-Random” method to query AF-Cluster has confounding factors: for instance, the “max_msa” random sampling that it relies on also performs clustering internally at the AF2 MSA processing step, so it is not a valid control for AF-Cluster.

To directly probe the role of evolutionary couplings in AF-Cluster, we performed a common control test that can be applied to any computational method where an MSA is used as input [23, 24]. We kept the first sequence of the MSA (the query sequence) the same, and shuffled residues within each column of the input MSA clusters, which ablates any potential evolutionary relationships beyond single-site conservation (**Figure 4A**). We compared the resulting structure predictions to those from the unperturbed MSA cluster. Performing this test using KaiB^TV^-4, first questioned in [2], shows that the MSA cluster resulted in substantially lower RMSD after zero recycles, whereas with shuffled MSAs, AF2 struggled to find the correct structure even after 12 recycles (**Figure 4B**). Having thoroughly investigated KaiB in [7], we wanted to further investigate clusters here that originally predicted both states for the other two proteins, RfaH and Mad2 in [7]. We used all previously-reported MSA clusters from [7] with more than 10 sequences as input to AF2, and ran with 4 random seeds, dropout, and all five AF2 models to test how consistent structure models were from different sequence clusters with increased stochasticity.

**Figure 4.**
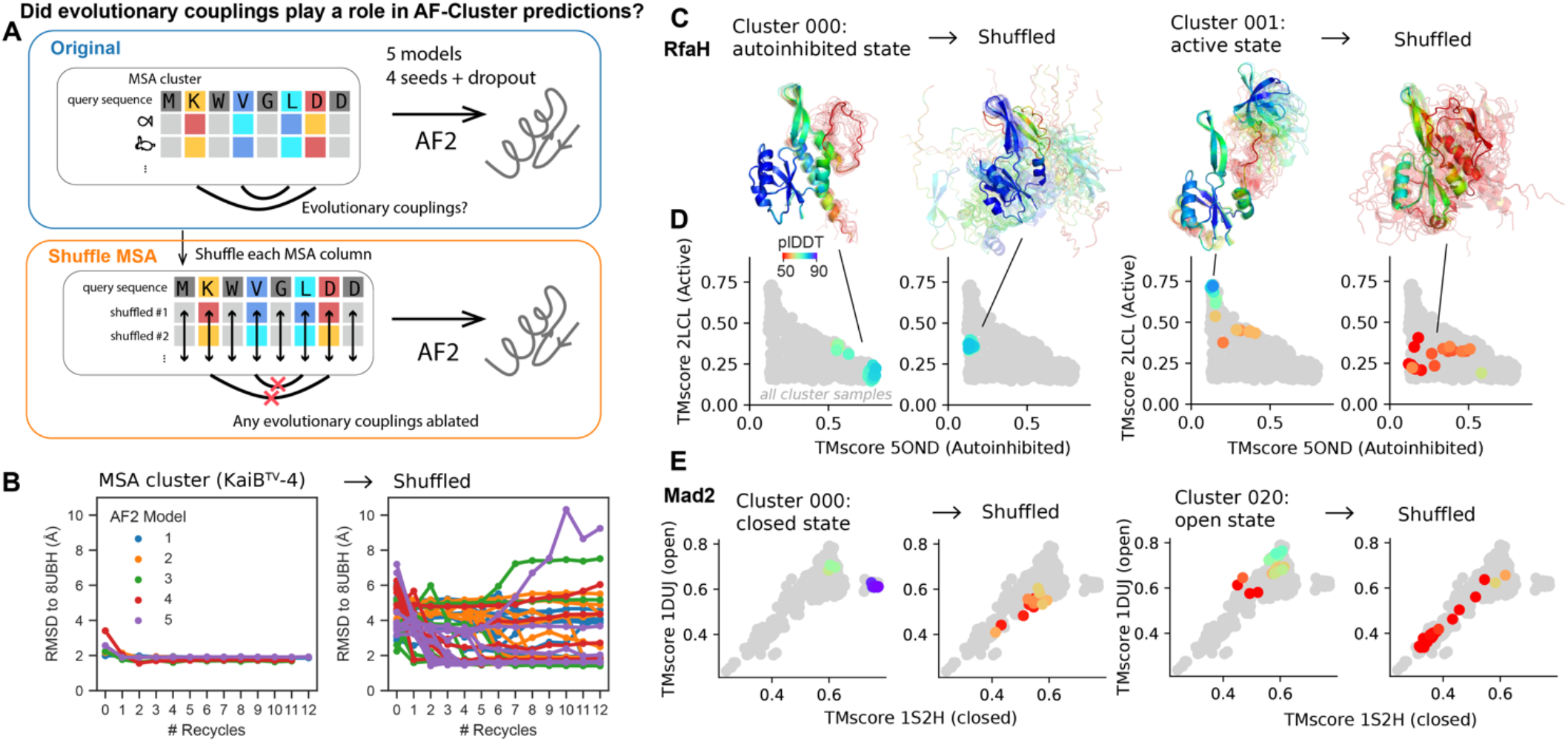
A) Scheme for testing if coevolution plays a role in AF2 predictions. B) Comparing AF2 predictions for MSA cluster for KaiB^TV-^4, the subject of investigation in [2], shows that coevolutionary information significantly aids AF2 in finding the known experimental structure. C) All models from stochastic sampling of two sequence clusters for RfaH from [7]. The AF-cluster predictions return the known structures, after shuffling no meaningful structures are obtained. D) Visualizing distribution of structures via TM-score to known states before and after MSA shuffling. Grey points in D, E depict all samples from AF-Cluster. E) For Mad2 as well, clusters that resulted in predictions of known states were hampered by shuffling columns to ablate coevolution.

**Figure 4C** depicts all structures from two representative MSA clusters that predicted the RfaH autoinhibited and active state. On the left are results obtained with 3 recycles without shuffling (left), and on the right are results obtained with shuffling (right) at 3 recycles. Even with dropout, which increases stochasticity in AF2, models from the same sequence cluster tended to go to the same part of structure space, as characterized by TM-score to two known structures of the fold-switching families (**Figure 4D,E**). In contrast, when evolutionary couplings are disrupted by shuffling, the predicted structures do not match the structure the original MSA cluster predicted. Equivalent figures for all clusters from [7] with 10 or more sequences are depicted in Appendix D.

We found that even with dropout, which can be interpreted as approximate sampling from the posterior distribution [25] and thus captures model uncertainty, the majority of the clusters that we originally reported as predicting one or the other state for RfaH and Mad2 still reported >50% of samples in the same state (see Methods and Appendix D). **Figure 5** depicts TM-score to the final state predicted from the original MSA cluster for both the original and shuffled MSAs, from 0 to 3 recycles. Note this is not dependent on pLDDT or other model quality metrics. This test reinforces that AF-Cluster is leveraging local co-evolutionary signals and sequence preferences to make its predictions. We note that only one of all these clusters showed equivalent ability to predict the RfaH autoinhibited state: cluster 049, which is the single cluster that was the subject of the majority of Schafer et al.’s investigations in Supplemental Figure 1 of [6].

**Figure 5.**
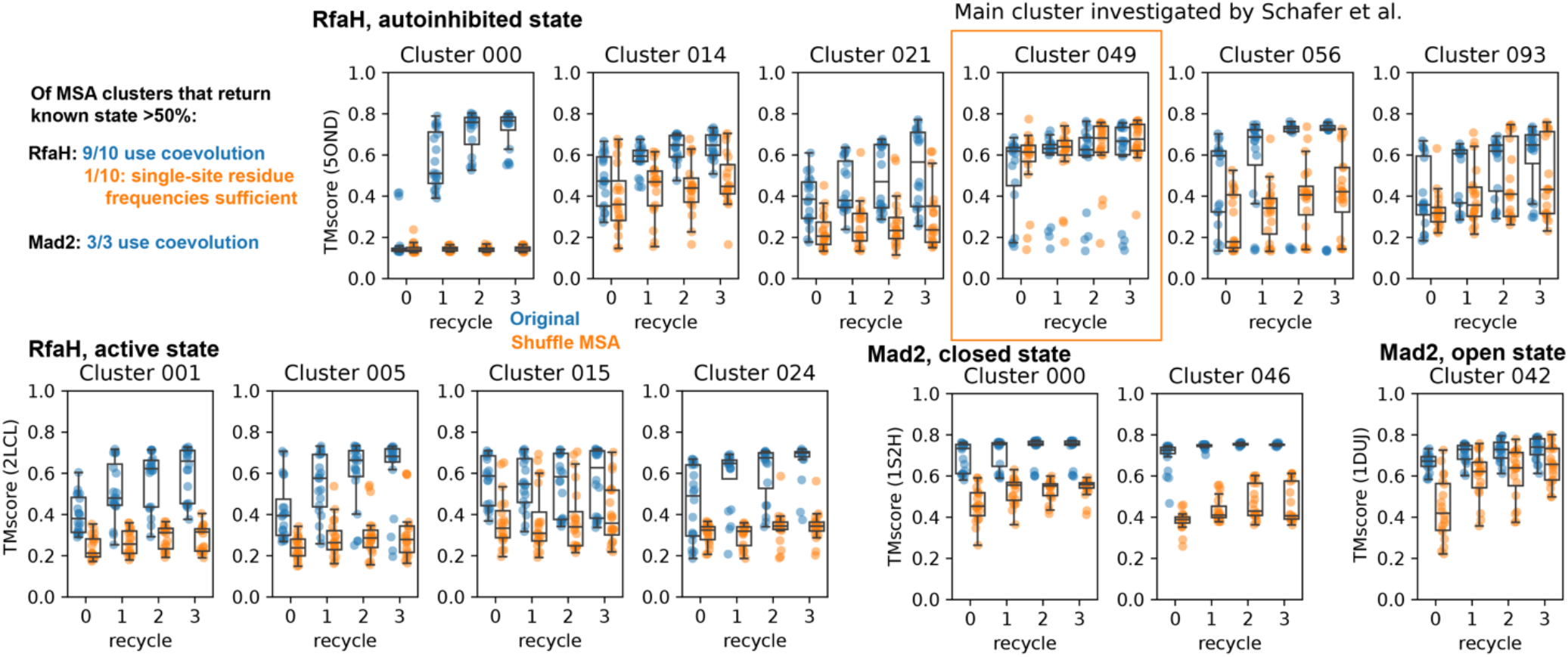
We analyzed all reported clusters for RfaH and Mad2 in [7] with increased stochasticity and 20 times more sampling than in [7]. We found that for the 13 clusters across RfaH or Mad2 that returned more than 50% of their samples as the same state after adding more stochasticity, all but one suffered when coevolution was ablated. This one was cluster 049 for RfaH, which was also the subject of the majority of Schafer et al.’s investigation.

One alternative hypothesis for why column shuffling might lead to reduced performance is that shuffling columns might be creating sequences dissimilar to ones AF2 would have seen during training, or disrupting memorized sequences that AF2 requires to predict these states. To test the viability of this alternative hypothesis, we used ProteinMPNN [26] to design 14-15 de novo sequences for KaiB, RfaH, and Mad2 folds that are dissimilar to any sequences in SwissProt as of June 2025, and used these as input MSAs to AF2 for the two experimentally-determined structures of KaiB, RfaH, and Mad2. We found that with these MSAs of de novo proteins, AF2 was able to robustly predict the structures (Appendix D), indicating that MSAs of natural proteins are not a requirement for AF2 to predict the metamorphic protein structures it is capable of predicting. If AF2 required memorized natural sequences, one would not expect these predictions to succeed.

## Conclusion

Comparing the main claims of [2] and [5], we want to highlight an intrinsic contradiction by the Porter lab in their own claims: the claim in [2] (single sequence is sufficient) to the second claim that CF-Random (which would be using a collection of sequences) is the way to predict the correct conformations [5].

The topic of how to predict multiple conformations from sequence is clearly far from solved. Experimental tests are critical, and are the major rate-limiter of methodological advancement. We therefore find this accusation in [5] to be disturbing: “Wayment-Steele et al. report a set of three mutations correctly predicted to switch the conformational balance of *R. sphaeroides* KaiB. Two of these three mutations caused AF2 to predict the same fold switch with pLDDT=68.15. Curiously, Wayment-Steele et al. do not report experimental tests of this double mutant prediction, leaving us to question its accuracy.”

One of [7]’s most impactful results was our experimental testing of computational predictions. We elected to make the triple and not the double mutant because of the higher pLDDT for the triple mutant, in detail explained and documented in [7] (c.f. Extended data Figure 6 in [7]). Strikingly, we made one single protein, and our NMR experiments fully verified our computational predictions. We note that such one to one agreement between prediction and experiment is rare, often in protein design and prediction many constructs are tested and only a few show such agreement.

## Methods

We have updated the public AF-Cluster repository to include exact commands to reproduce every model prediction in our original paper [7], as well as models presented here, at https://github.com/HWaymentSteele/AF_Cluster/blob/main/complete_methods.md.

To generate the data in Figure 1c [6],d of this preprint, we ran ‘run_af2.py’ available in the github repository of [7], using either the local-10 MSA corresponding to KaiB^TV^-4 from [7], or the sequence of KaiBTV-4 as a single sequence, varying the model number and number of recycles.

To generate the data in Figure 2d of this preprint, we followed the CF-random methodology described in the supplemental information in [5]. To summarize, we ran ColabFold for KaiB^TE^ (sequence in 2QKE) with 33 random seeds and otherwise default settings, and ran ColabFold for KaiB^TE^ with 33 random seeds and max_msa:extra_msa=1:2 and otherwise default settings.

To generate the data in Appendix 1, we ran AF2 with and without masking, with old parameter versions and new parameter versions, and using either the complete RfaH MSA reported in [7] or cluster 49 from [7]. Code is available in https://github.com/HWaymentSteele/controls_04feb2024.

To generate the data presented in Fig. 3 for resampling the MSA cluster, we ran ColabFold using the MSA cluster for KaiB^RS^ with 5 random seeds, 5 models, and 3 recycles. To re-create CF-Random sampling, we ran ColabFold for KaiB^RS^ with 5 random seeds, 5 models, 3 recycles, and max_msa:max_extra_msa = 1:2.

To generate the runs presented in Fig. 4B, we ran ‘run_af2.py’ available in the Github repository corresponding to [7]. For shuffled replicates, we ran 10 replicates, where <REP> is between 0 and 9: python run_af2.py data_sep2022/06_kaibtv4_followup/msas/scrambled/scramble_<REP>.a3m -- model_num <NUM> --recycles <N_RECYCLES> --output_dir.

To generate the runs presented in Fig. 4D,E, and Appendix D, we ran ‘coevolutionary_ablation_2025/run_all.py’ available in the Github repository corresponding to [7]. The same runs can be generated using the ColabFold environment present in the AF-Cluster Colab notebook presented here.

For Figure 5, we ran all MSA clusters from ref. [7] that either were the MSA clusters that generated the structure models for RfaH and Mad2 originally depicted in ref. [7], as well as any MSA cluster with 10 or more sequences. This resulted in 49 MSA clusters in total for RfaH and 32 in total for Mad2. For each of these, we ran these in all 5 models, with dropout, for 3 different seeds, and for 3 recycles.

We categorized RfaH structure outputs as autoinhibited or active states based on TMscore of the C-terminal domain, using a TMscore cutoff of 0.65 for both states. This was based on inspecting returned models. We categorized Mad2 structure outputs as open or closed state using a TMscore cutoff of 0.72. These TMscore cutoffs were based on inspecting resulting models.

Figures D1,2 in Appendix D depict all clusters and their shuffled counterparts at 3 recycles for RfaH and Mad2, respectively. Figure D3 depicts the number of clusters that returned either state, either original MSA cluster or shuffled cluster, at different fraction cutoffs of all samples generated from the MSA.

ProteinMPNN design. We used the ProteinMPNN [26] implementation on HuggingFace (https://huggingface.co/spaces/simonduerr/ProteinMPNN) to design sequences corresponding to the structures 2QKEE, 5JYTA, 6C6SD, 1S2HA, 1DUJA. Because the autoinhibited state of RfaH (5ONDA) has experimentally-unresolved residues, we used the structure from cluster 049 from the AF-Cluster data respository [7]. For all targets, we used a temperature of 1.0 and generated 15 sequences. We used BLAST[27] to compare each of these sequence sets to both the PDB and SwissProt on June 15, 2025 and removed any sequences that resulted in hits to either database. This removed 1 sequence each for 2QKEE, 5JYTA, and 5ONDA. No further statistically significant hits at BLAST’s default settings were identified. These sequence sets were then used as MSA inputs to ColabFold[19]. The structures depicted in Appendix D4 are the top-ranked structure predictions from ColabFold by pLDDT.

## Data availability

All data corresponding to new calculations presented here are at https://github.com/HWaymentSteele/controls_04feb2024 or at https://github.com/HWaymentSteele/AFCluster/coevolutionary_ablation_2025.

## Code availability

Code to reproduce the new calculations presented in this response are available at https://github.com/HWaymentSteele/controls_04feb2024 or at https://github.com/HWaymentSteele/AFCluster/coevolutionary_ablation_2025.

An implementation of AF-Cluster in ColabDesign, which allows users to more easily integrate the AF-Cluster DBSCAN-based clustering step with other AF2 sampling methods, is available at https://github.com/HWaymentSteele/AF_Cluster/blob/main/AF_cluster_in_colabdesign.ipynb.

## Acknowledgment

We thank Gina El Nesr and members of the Ovchinnikov lab for useful discussions. This research was funded by the Jane Coffin Childs foundation (H.K.W-S.) and Howard Hughes Medical Institute (D.K.). S.O. was supported by NIH [DP5OD026389] and NSF [MCB2032259].

**Figure A1.**
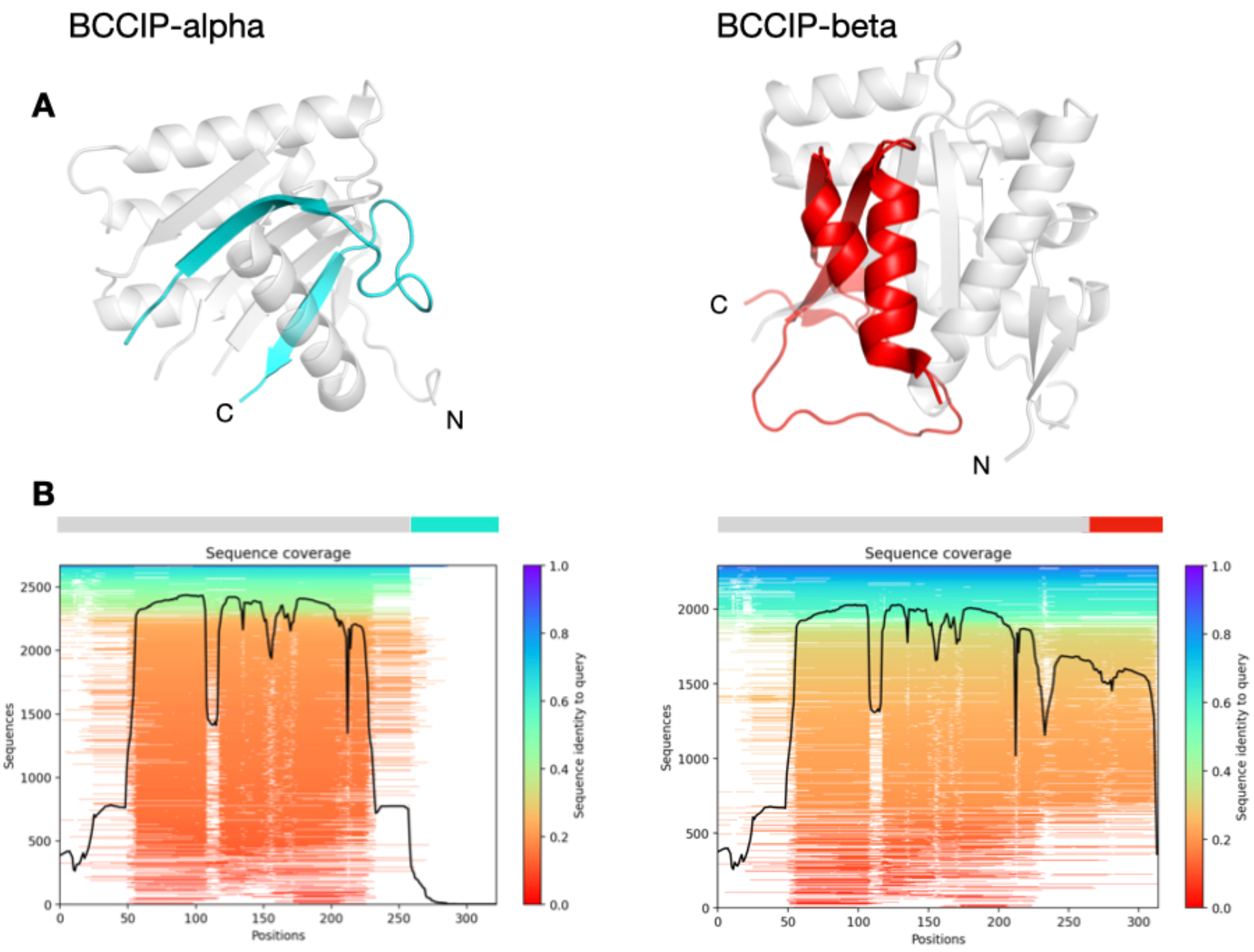
Appendix A. No sequence coverage for fold-switching region in BCCIP-alpha. A) Structure of BCCIP-alpha (left, PDB: 8EXE:B) and BCCIP-beta (right, PDB: 7KYS:A). Common sequences are depicted in grey, differing sequences are depicted in cyan in BCCIP-alpha and in red in BCCIP-beta. B) Sequence coverage of BCCIP-alpha (left) and BCCIP-beta (right), generated in ColabFold. Sequence coverage of the alternatively spliced sequence in BCCIP-alpha is minimal (cyan).

## Appendix B “Sequence Clustering Confounds AlphaFold” as preprint uses different AF2 parameters and cannot be compared

Given the discrepancy between [5]’s model settings and ours, we investigated the extent to which different model settings matter for RfaH predictions. ColabFold implements AF2 internally and should return identical solutions assuming the same MSA input is used. Differences may arise if different model weights or settings are used. Our reported values use the same model (model_1_ptm) for both full and clustered MSAs. We confirmed that our reported pLDDT values are not significantly affected by two differences between default ColabFold and our AF2 script: random seeds and use of masking (see Appendix 1). However, we could only reproduce pLDDT values reported in [5] by using an older version of AF2 parameters that are not default in ColabFold, were not used in AF-Cluster, and significantly affect reported values (Appendix 1). Therefore, [5]’s calculations cannot be compared to what we reported in the paper.

The pLDDT values for both the full-MSA and the clustered sequences depend on the specific AF2 model, so it is essential that controls are performed with the same AF2 implementation and models. With proper controls in place, our benchmarking supports our original finding, that clustering the input MSA and using these clusters as input achieves higher pLDDT for the RfaH autoinhibited state than the full MSA, and that the claim made by [5] is false. Again, we want to emphasize we do not expect all clusters to have higher pLDDT.

Figure B1 depicts structure models of a random protein with low MSA coverage generated with the DeepMind AlphaFold notebook and the ColabFold notebook using the same MSA and settings, demonstrating that the resulting structure predictions are the same.

**Figure B1.**
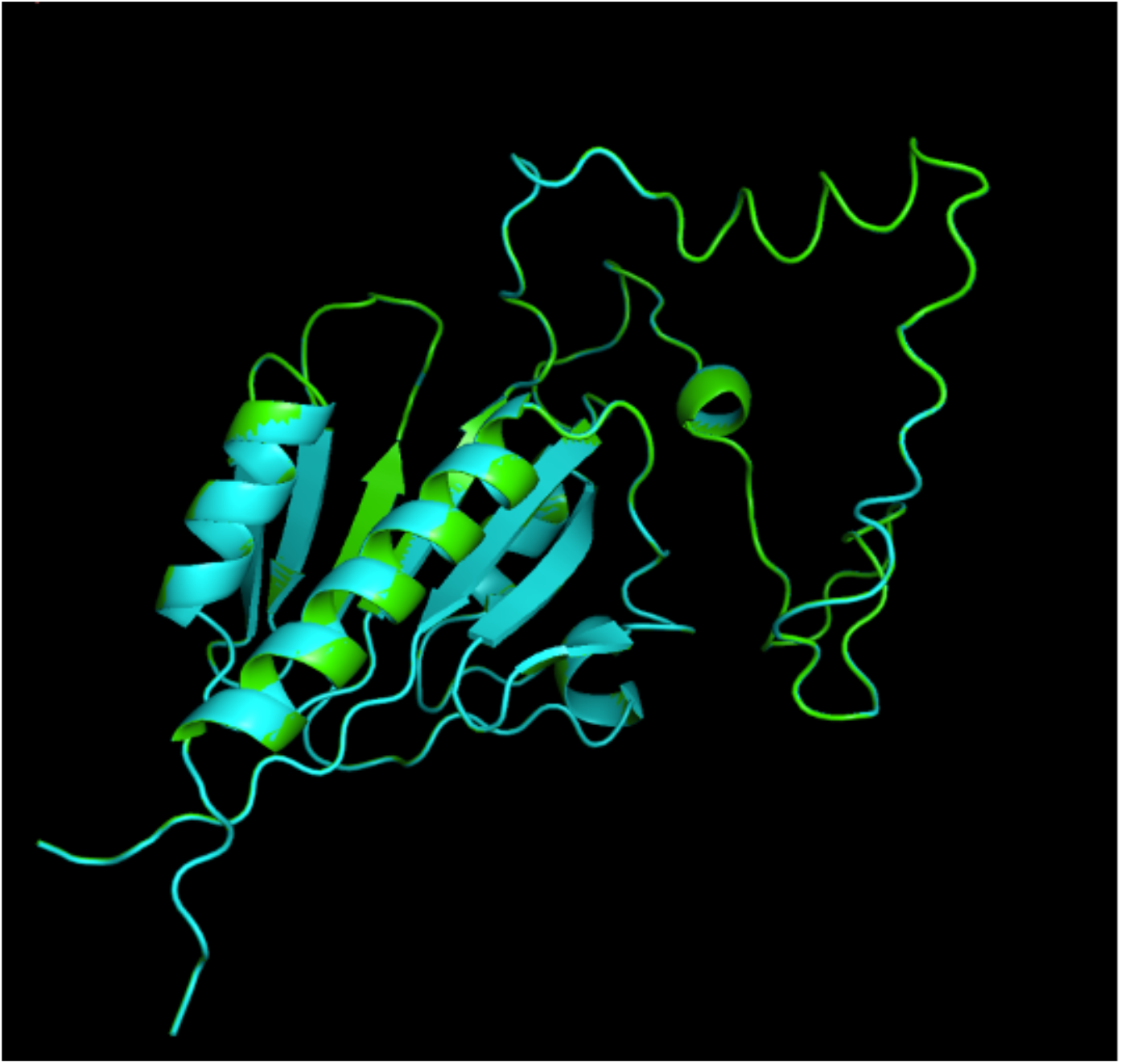
AlphaFold and ColabFold return the same result when given the same MSA and same settings. Green: Deepmind AlphaFold notebook. Cyan: ColabFold AlphaFold notebook (use_bfloat16=False). The outputs are identical and superimpose, without need to align.

Crucially, the only settings that allowed us to reproduce [5]’s RfaH calculations from a full MSA was by using older versions of AF2 parameters, named with the convention ‘model_[1,2,3,4,5]’. The current default parameters in ColabFold, and those used in AF-Cluster, are a newer set of parameters from a more recent training of AF2. These are named according to the convention ‘model_[1,2,3,4,5]_ptm’. Figure B2 depicts pLDDTs of RfaH structures sampled with 50 random seeds in all 5 parameter sets, comparing the old and new parameter sets. This demonstrates that 1) the old and new parameters result in significantly different pLDDT values, and 2) between parameter sets, also termed AF2 models, the results can change quite significantly. Therefore, controls varying other aspects must be performed using the same parameter set, as we did in our paper, but was not done in [5].

**Figure B2.**
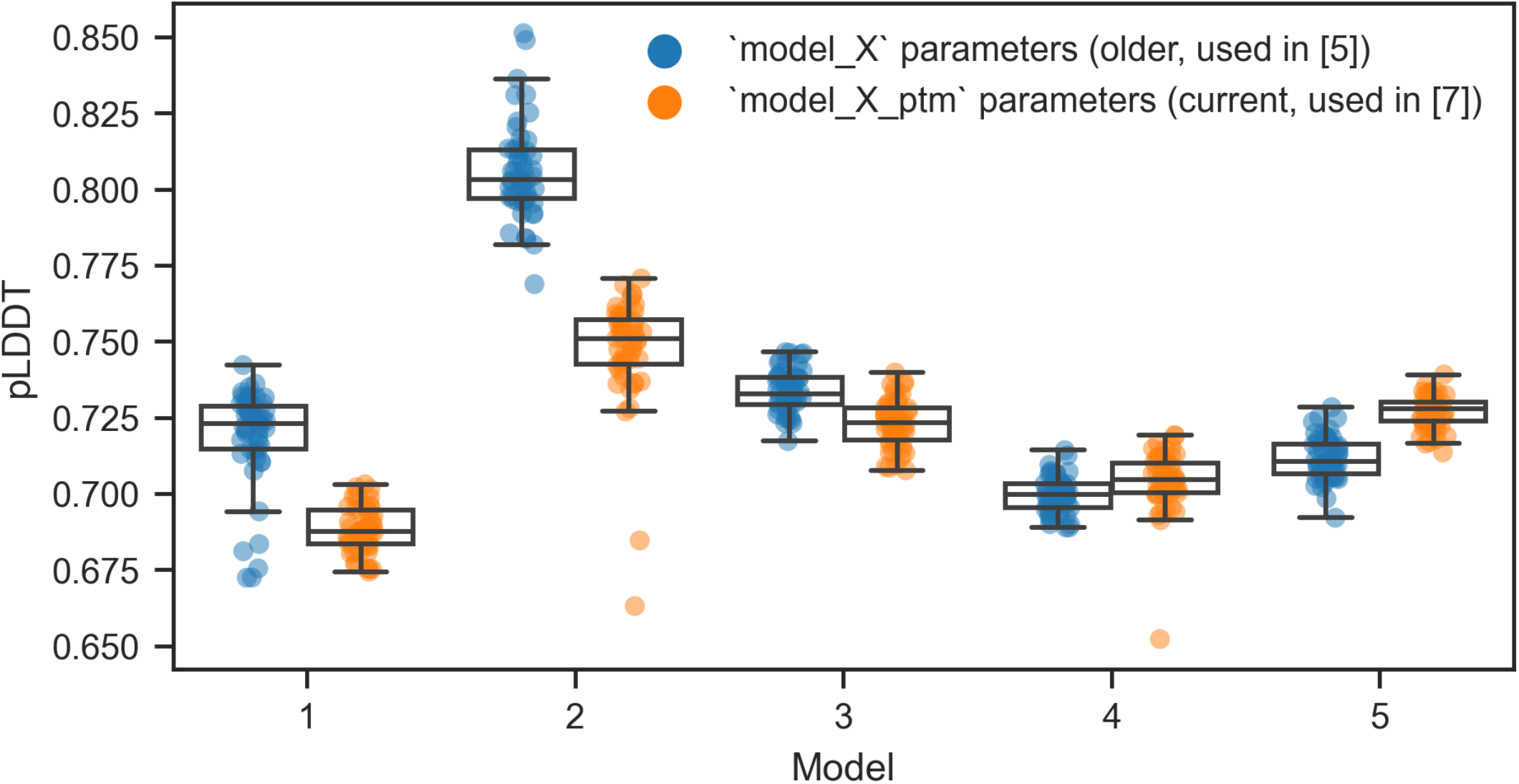
pLDDTs of RfaH structures vary significantly based on parameter version and parameter model number. Box plots depict median and 25/75% interquartile range, whiskers = 1.5*interquartile range.

One discrepancy we realized in our paper is that the RfaH full MSA predictions (Extended Data Figure 6a in ref. [7]), run with ColabFold, included random masking, whereas the models generated with ‘run_af2.py’ in the rest of the paper did not include random masking. We repeated the same control without random masking, and found the results to be very similar within model 1, which is what we reported to compare to AF-Cluster, as all AF-Cluster runs were only performed in model 1. In 4 of 5 AF2 models (each except model 4), we find this to be the case for the same cluster investigated in [5] (Figure B3). Therefore, the claims made by [5] are incorrect.

**Figure B3.**
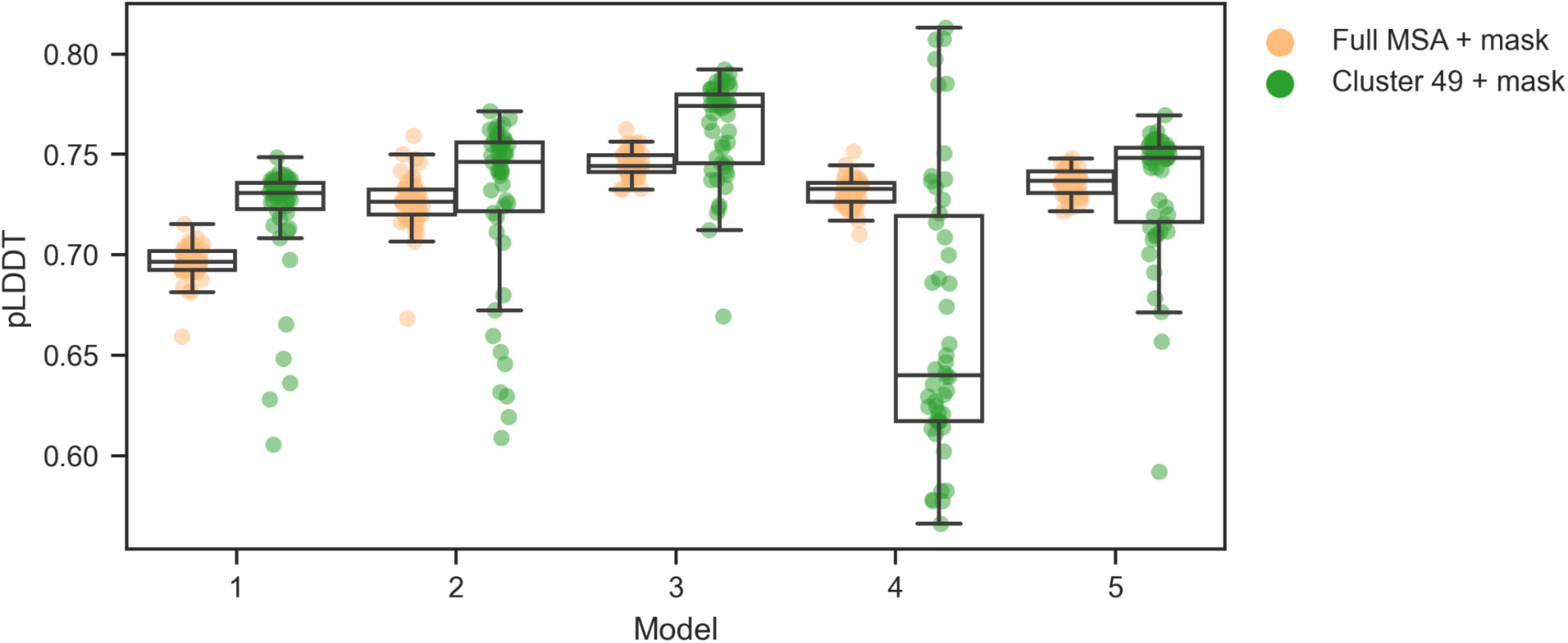
Comparing models from the example MSA cluster 49, used in [5], with correctly-compared settings (masking, current parameter version) indicates that in 4 of 5 AF2 models, the pLDDT from MSA cluster 49 (green) is higher than the pLDDT from the full MSA (orange). Box plots depict median and 25/75% interquartile range, whiskers = 1.5*interquartile range.

## Appendix C.

[5] provides insufficient evidence to substantiate their claim: “To the contrary, no amino acid contacts unique to the autoinhibited a-helical form were observed, but weak contacts unique to the active b-sheet form were present (Supplementary Figure 1b).” We have boxed this feature in red in Figure C1.

The claim of “weak contacts unique to active b-sheet” (red box) cannot be validated since this region is not resolved in the crystal structure of the RfaH autoinhibited state (PDB: 5OND). In fact, contacts within those regions are formed in both the active (in red, Figure C1F,H) and inactive state (in red, Figure C1E,G), and thus cannot be claimed to be unique to active state b-sheet.

**Figure C1.**
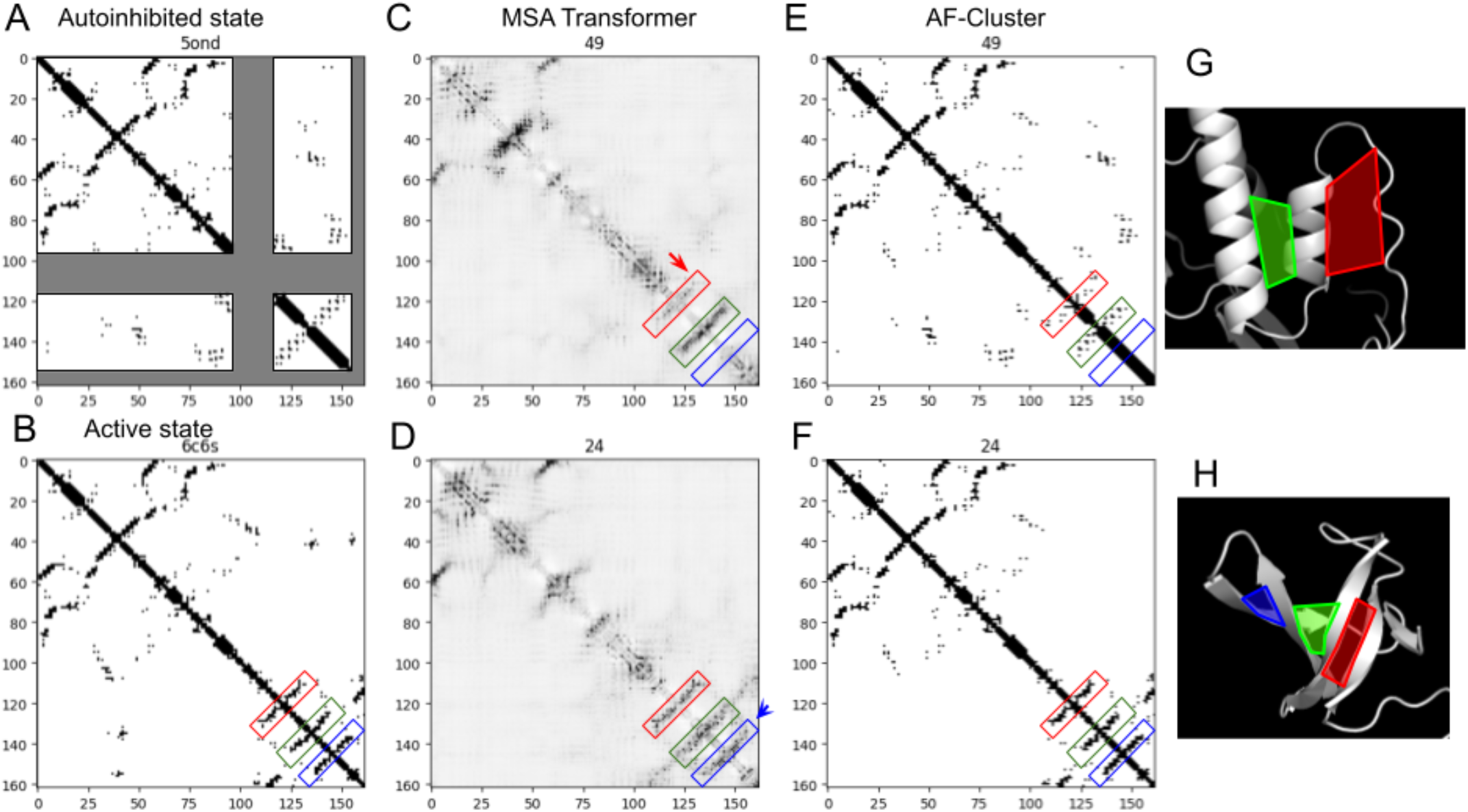
Comparing contacts in the experimentally-determined structures of the autoinhibited state (A) and the active state (B) of RfaH to contact predictions from MSA Transformer (C,D) and AF2 predictions (E,F,G,H) from MSA clusters. Schafer et al. [5] claim that the contacts identified in cluster 49 by MSA transformer are unique to the beta-strand of the active state (C, red arrow). However, this region is absent from the crystal structure of the autoinhibited state (gray area in A). These red boxed contacts are also present in the AF-Cluster predicted model of the autoinhibited state (red box in E, G), so they are not unique to the beta-strand as Schafer et al. claim. In contrast, the contacts that are unique to the active state (D, blue arrow) are not present in the MSA transformer contacts from cluster 49. However, they are observed in MSA Transformer prediction from cluster 24, which predicts the active state.

To assess this more systematically, we input all RfaH MSA clusters from ref. [7] into MSA Transformer (not just one, as presented in refs. 5, 6), and calculated the differences in contacts between the MSA clusters that produced the structure models for RfaH shown in Figure 4b of ref. [7] (5 for each cluster) and a baseline from across all clusters (Figure C2). We created this baseline by randomly selecting 5 clusters, taking their average, and repeating this 100 times and taking the average.

If we take the difference between these sets of clusters and the baseline, we see clear features that are enriched in each set of MSA clusters that correspond to the same state predicted by AF2 in the AF-Cluster method (Figure C2B). This clearly supports the idea that there are distinctive coevolutionary signals for each state that both MSA Transformer and AF2 are detecting.

**Figure C2.**
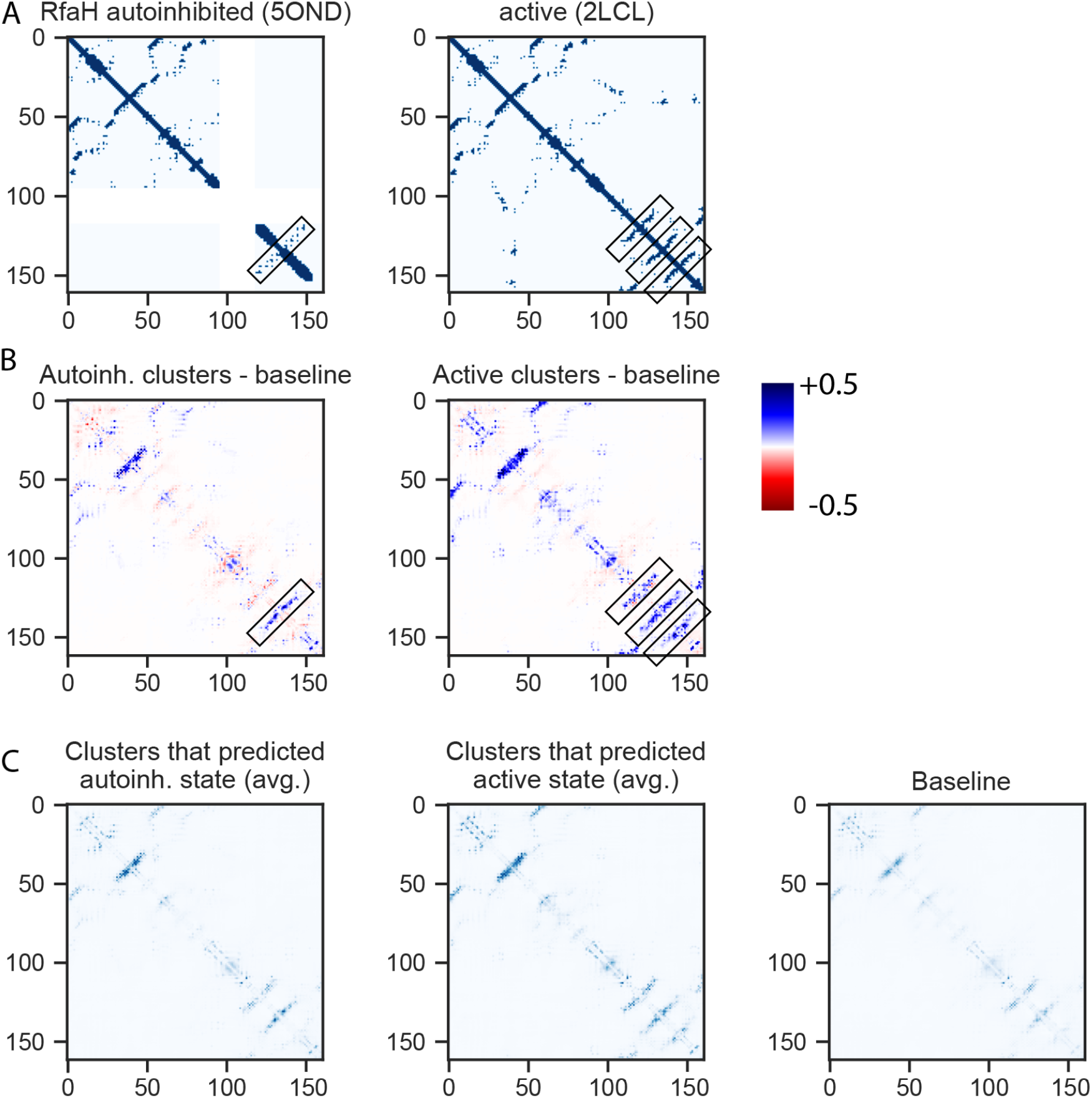
A) Contacts in experimentally-determined structures of RfaH autoinhibited state (5ONDA) and Active state (6C6SD). White areas correspond to residues that are not resolved in 5ONDA. Contacts unique to autoinhibited state and active state are boxed. B) Taking average of all clusters that predicted Autoinhibited state in ref. 7 and subtracting baseline contacts across all clusters clearly shows that the clusters that predicted the Autoinhibited state in AF-cluster also have signal in MSA Transformer for the autoinhibited state. The same is true for the active state (boxed features). C) Averaged raw outputs from clusters that predicted the autoinhibited and active state, and the baseline comparison: randomly select 5 clusters and average over 100 bootstrap iterations.

## Appendix D Supporting information for MSA column shuffling analysis.

**Figure D1.**
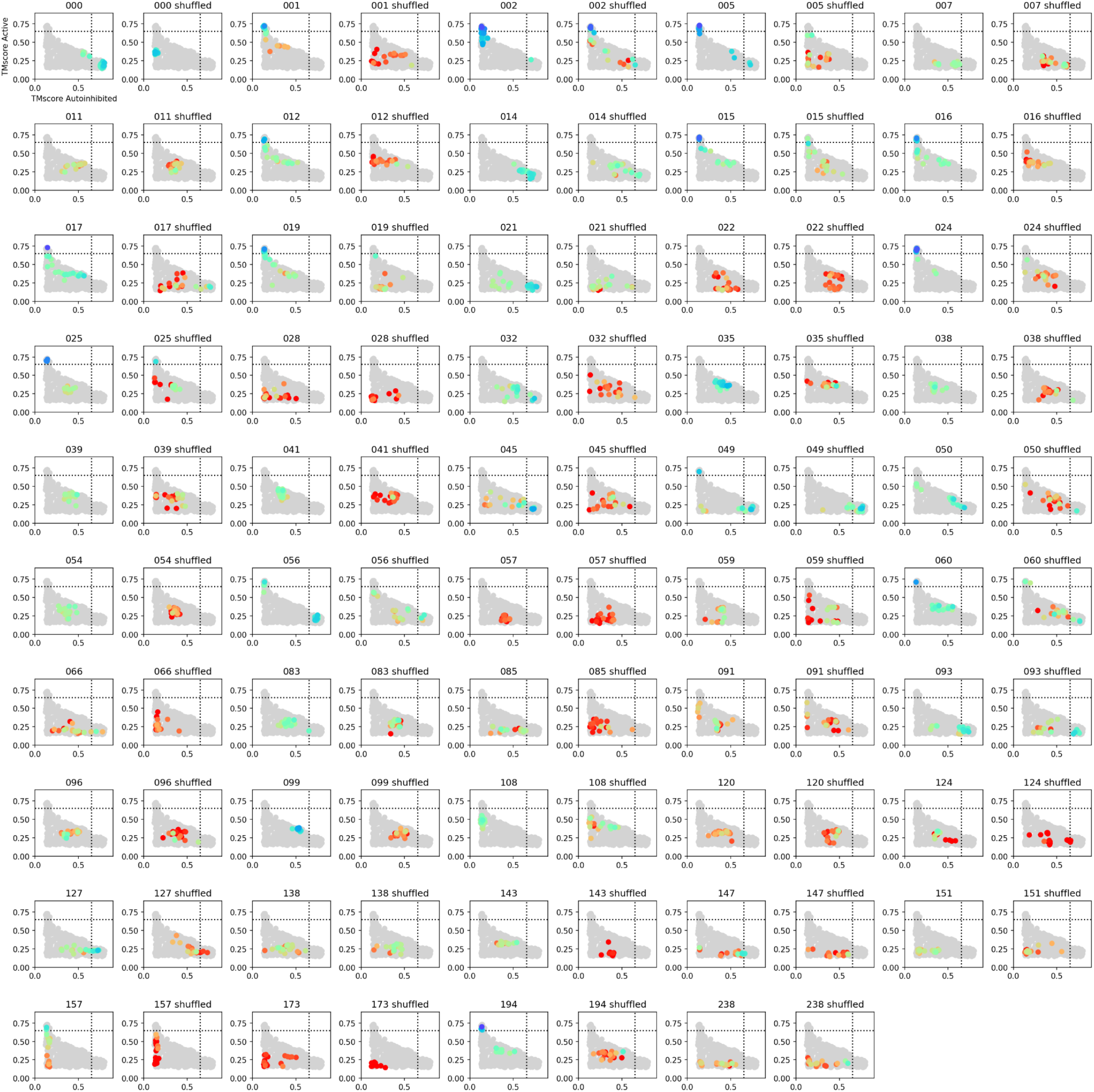
All RfaH clusters, original and shuffled. Grey: all structures sampled. Structures for each cluster are colored by pLDDT (red: 50, blue: 90).

**Figure D2.**
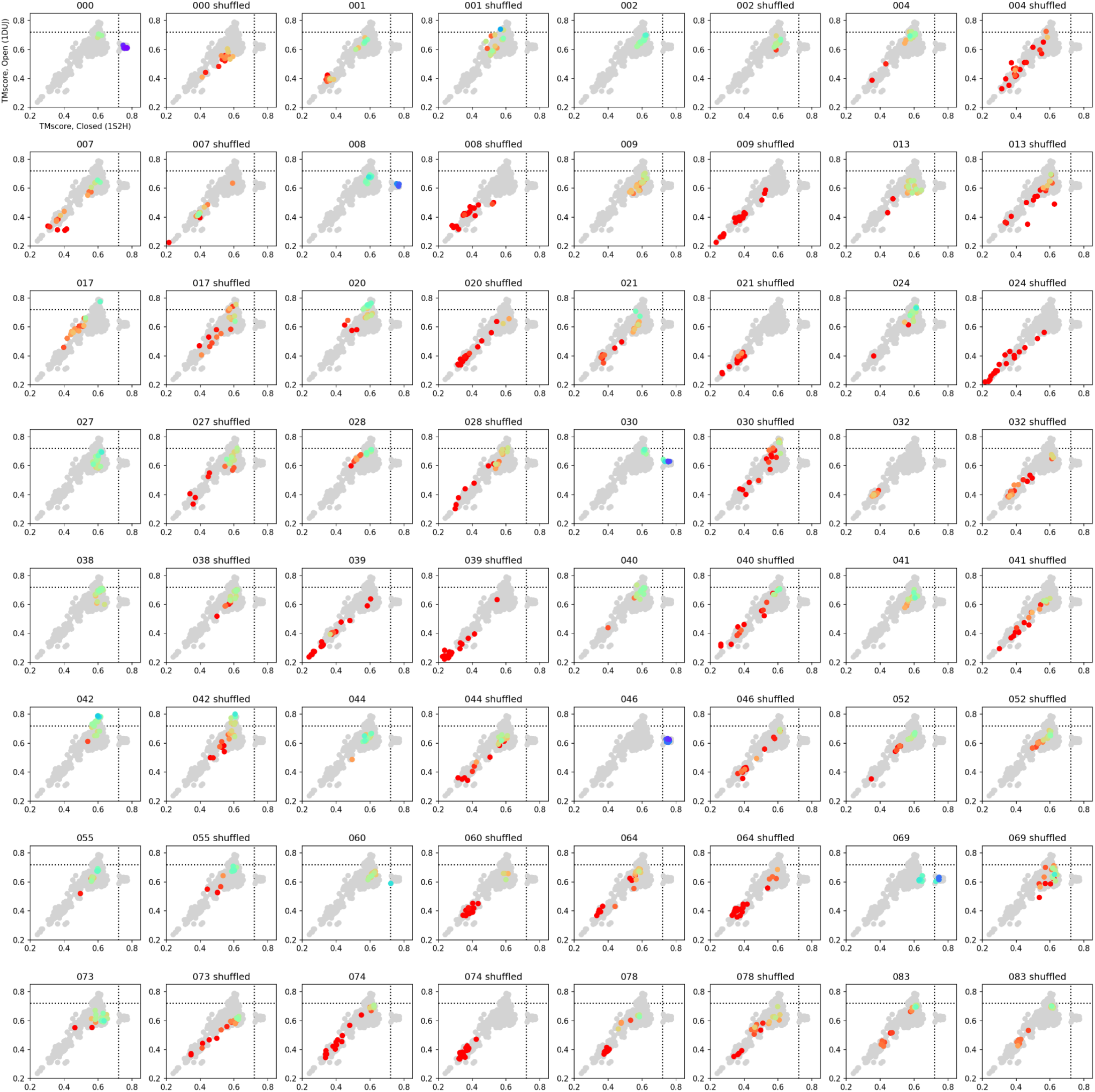
All Mad2 clusters, original and shuffled. Grey: all structures sampled. Structures for each cluster are colored by pLDDT (red: 50, blue: 90).

**Figure D3.**
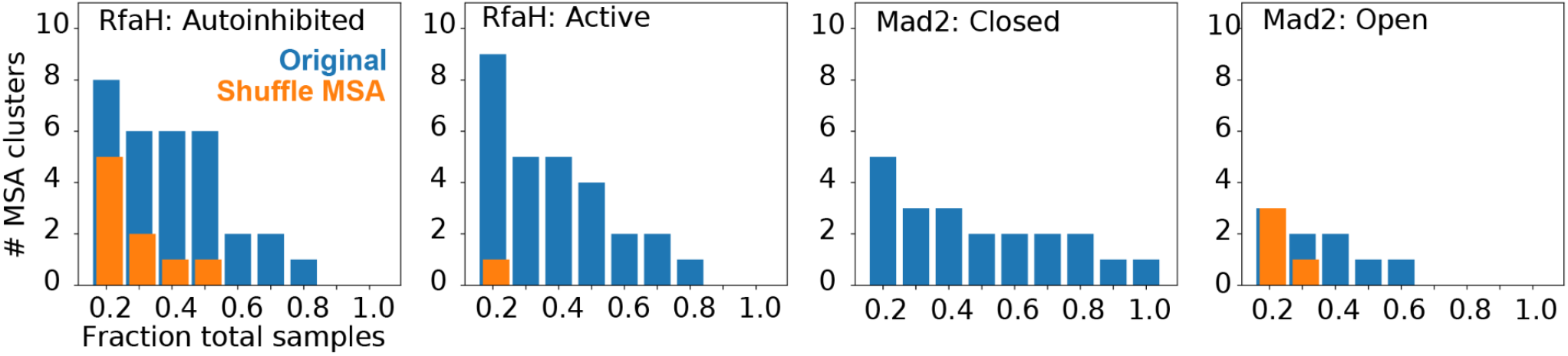
Comparing the number of MSA clusters that returned different states at different fractions of all samples, for original vs. shuffled MSAs. In Figure 4, we show clusters that return one or the other state with a fraction of 0.5 or more.

**Figure D4.**
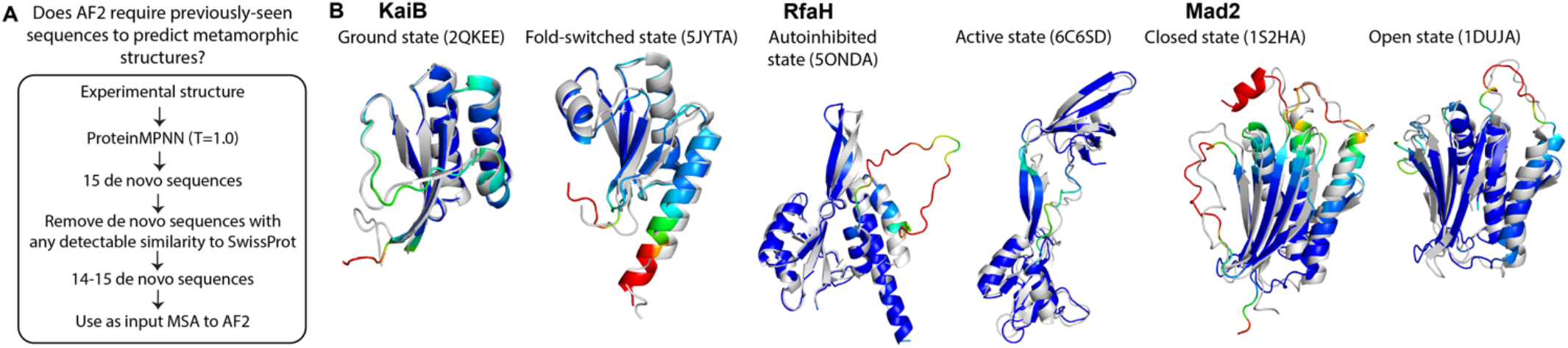
A. Workflow for generating de novo sequences for KaiB, RfaH and Mad2 folds with no significant hits in SwissProt to test if AF2 requires previously-seen sequences (memorization) for predicting structures of metamorphic proteins. B. Each experimental structure is in grey, top ranked-structure by pLDDT generated with ColabFold using de novo sequences is depicted in color (red: pLDDT=50, blue: pLDDT=90). Sequences are in Table 1.

---

**Table 1.**
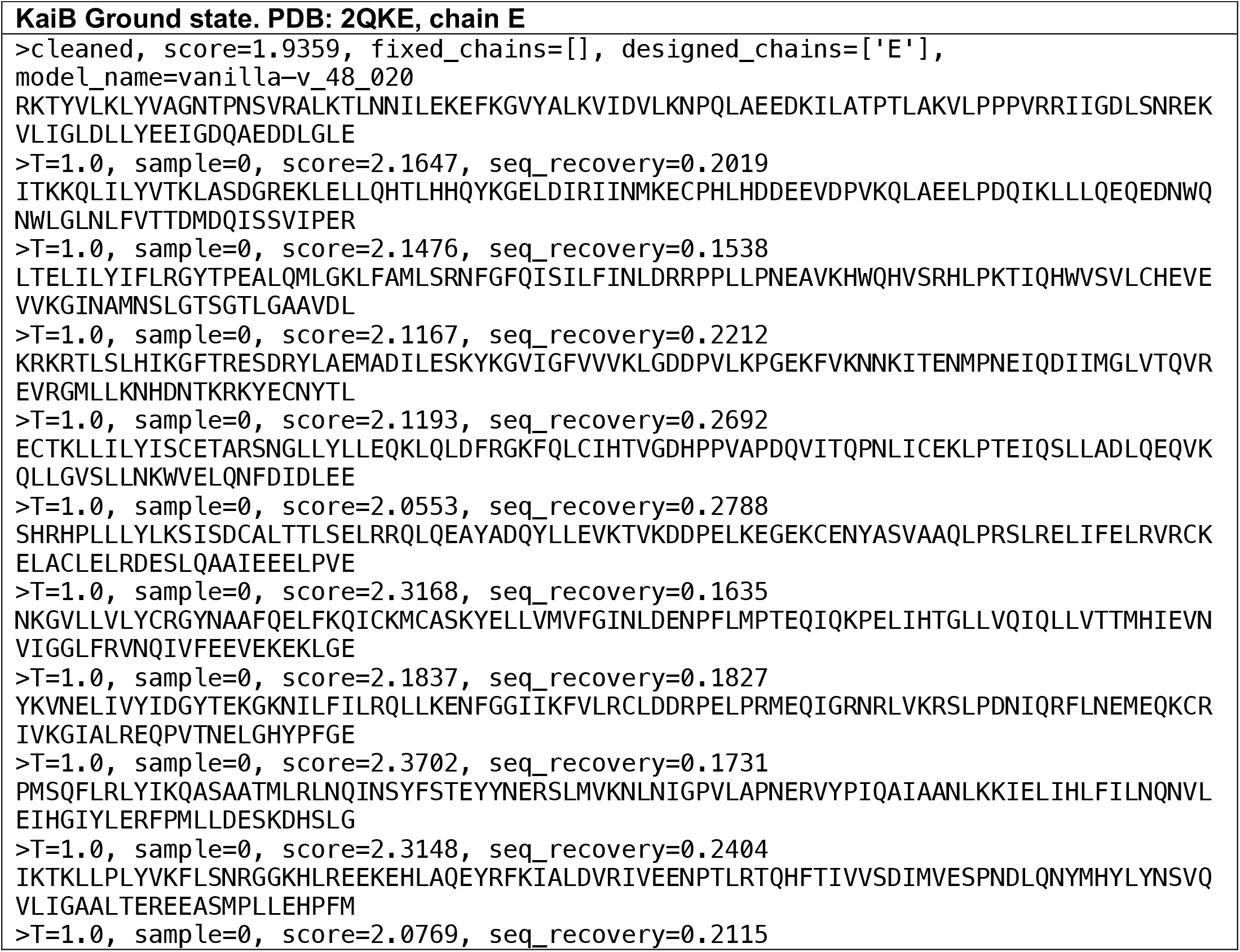
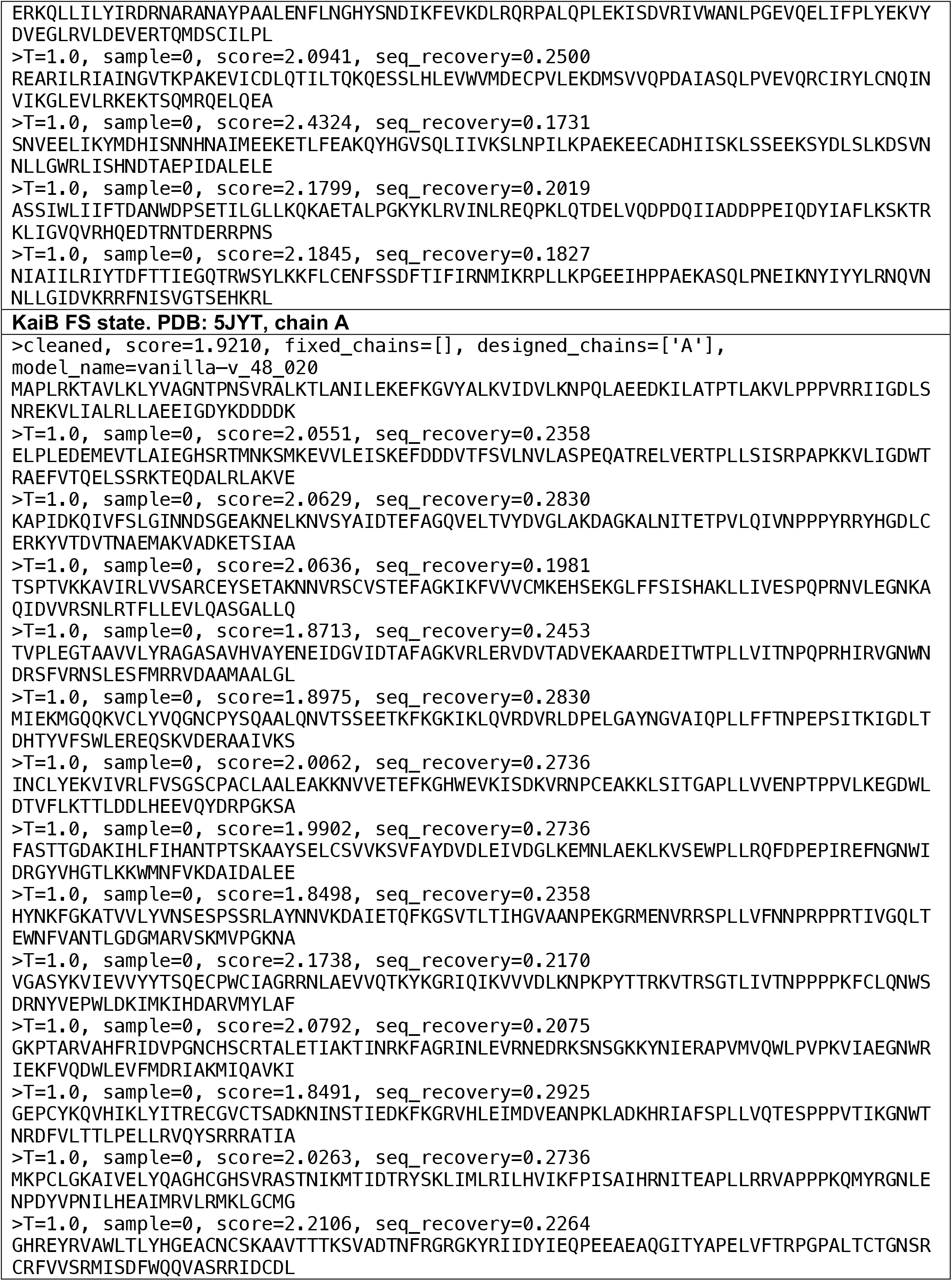
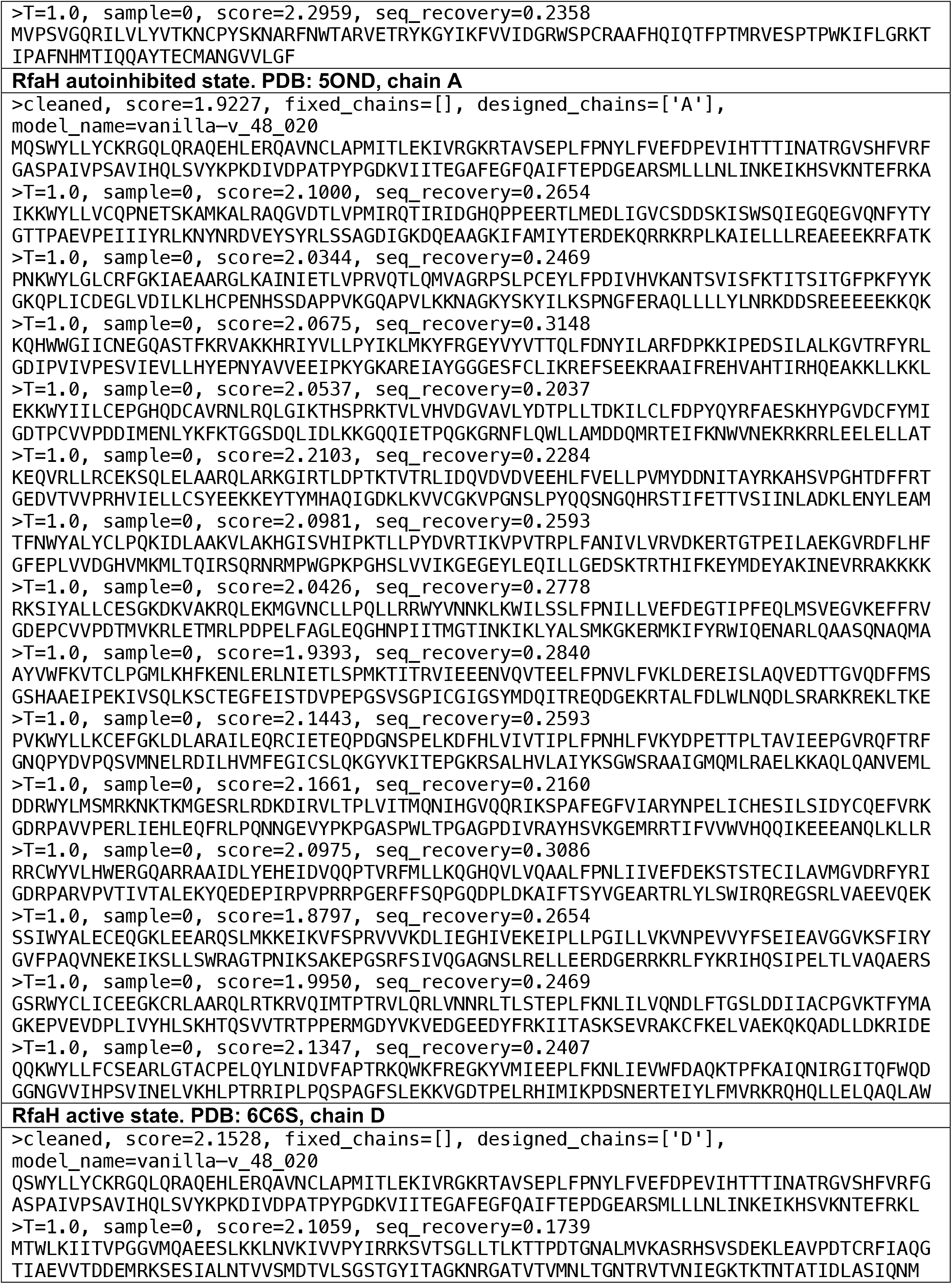
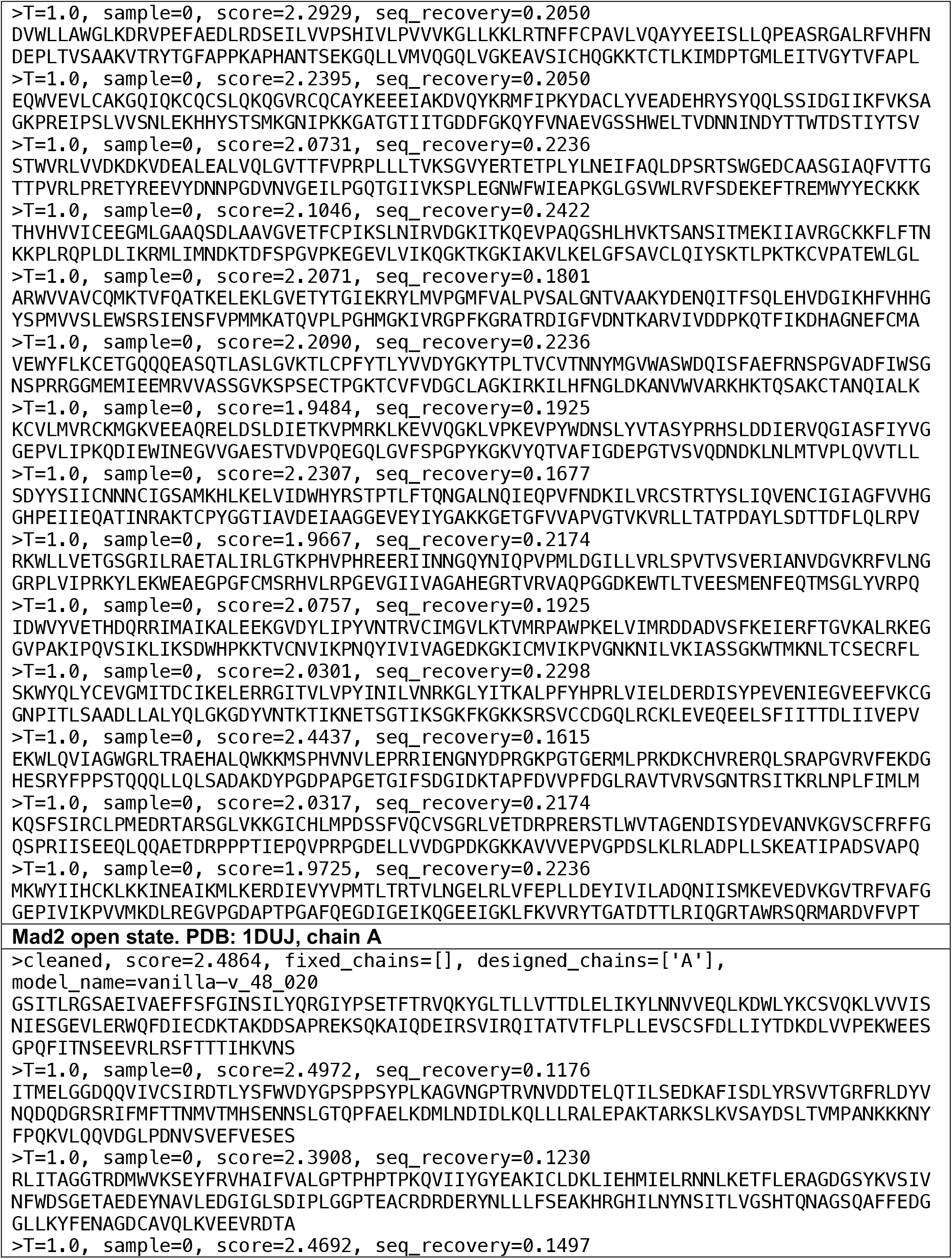
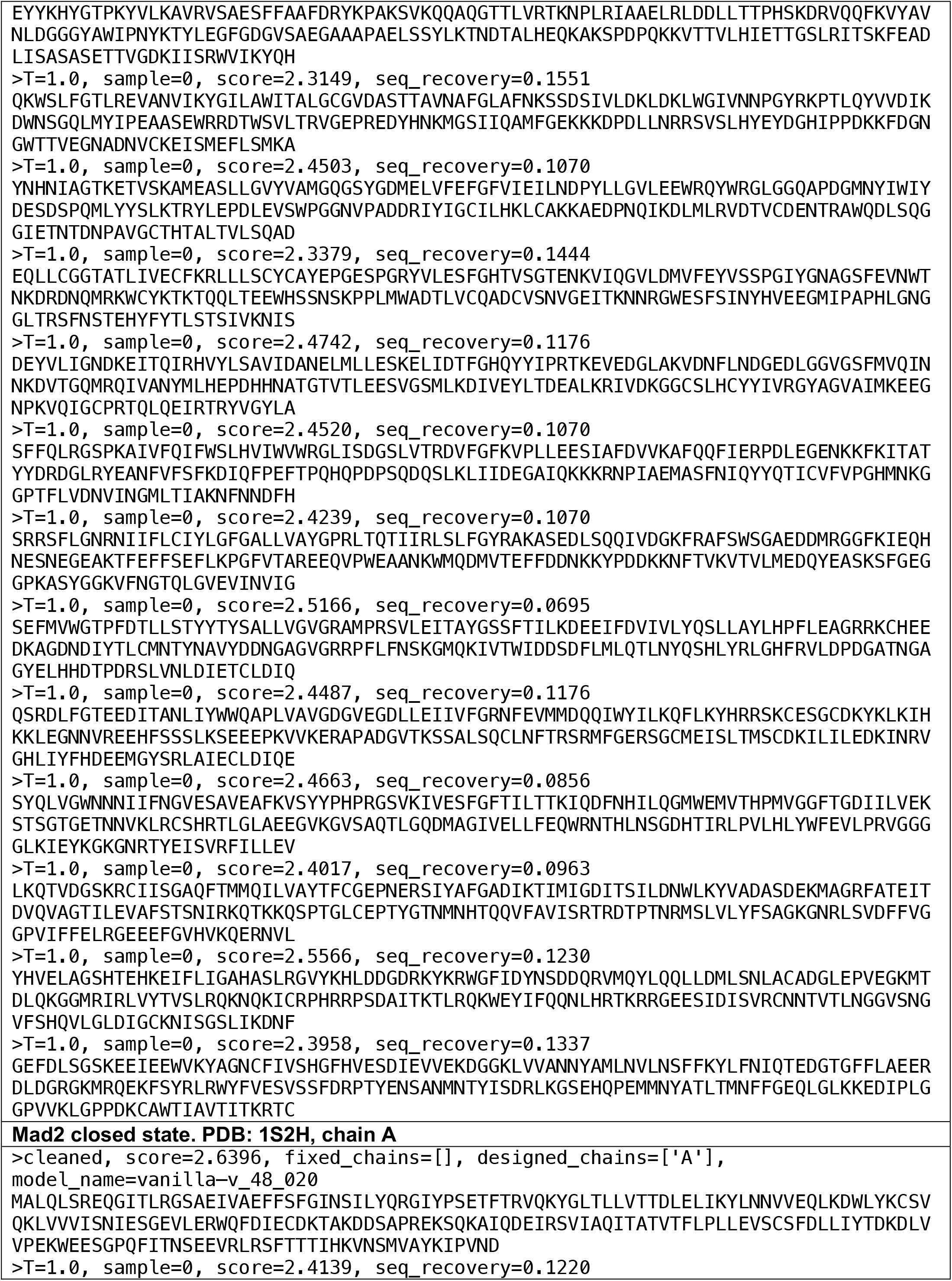
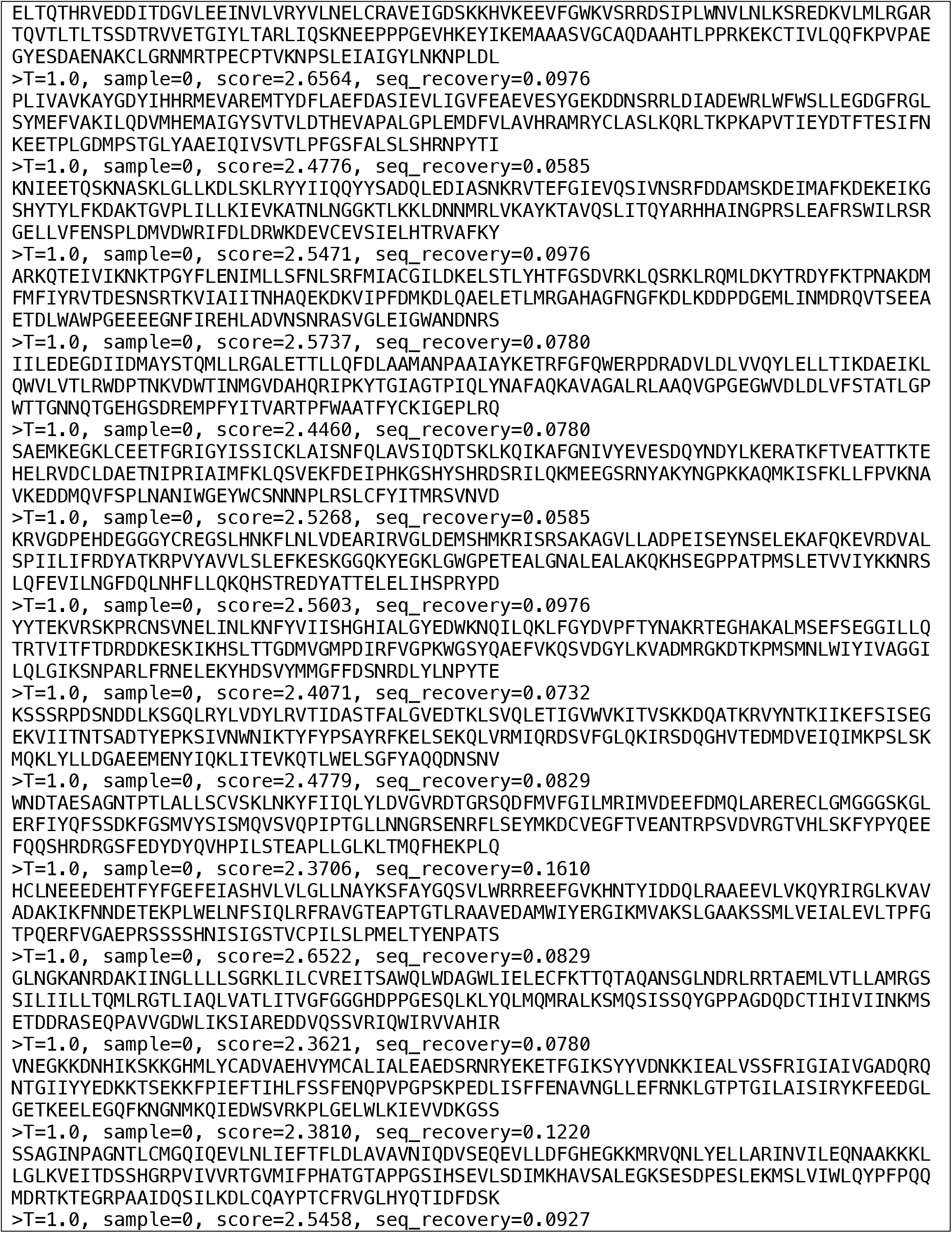
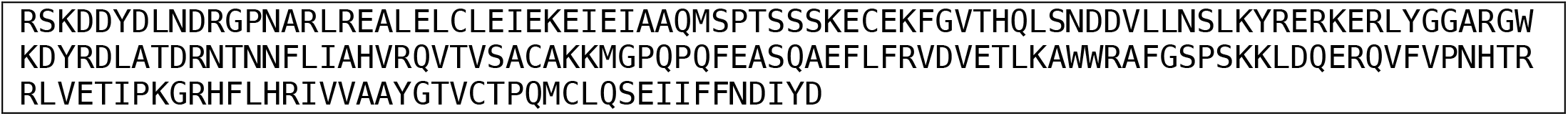
De novo fold switching sequences designed with ProteinMPNN.

